# Ca^2+^ and DRP1 drive endocytic lysosome reformation at tripartite contact sites

**DOI:** 10.64898/2026.01.30.702748

**Authors:** Samit Desai, Emma Martín-Sánchez, Danilo Ritz, Alexander Schmidt, Anne Spang

## Abstract

Lysosomes are essential in maintaining cellular health. Endocytic lysosome reformation (ELR) regenerates functional lysosomes following degradation of endocytic cargo, yet the mechanisms driving this process remain largely unknown. Here, we define the molecular machinery underlying ELR. We find that unlike autophagic lysosome reformation (ALR), ELR proceeds independently of mTOR and dynamin 2, but requires the mitochondrial fission GTPase DRP1. DRP1 mediates scission of endolysosomal tubules at contact sites with the endoplasmic reticulum (ER) and mitochondria. Disruption of DRP1 function or ER-endolysosome contact results in elongated tubules, indicating defective lysosome reformation. Moreover, mitochondrial activity is essential for tubule initiation, and Ca^2+^ transfer from endolysosomes to mitochondria is crucial for ELR onset. Our findings reveal a dual role for mitochondria in ELR: first in ELR initiation and second in DRP1-dependent tubule fission at ER-mitochondria-endolysosome tripartite contact sites, uncovering the previously unappreciated role of mitochondria in endolysosome remodeling and fission.

## Introduction

Endocytosis is a process by which cells internalize membrane proteins destined for either recycling or degradation. This process follows a well-defined route starting at the plasma membrane. Small vesicles carrying endocytosed cargo fuse together to form early endosomes^1,2^. Depending on the fate of the cargo, it is either sorted for recycling or destined for degradation. The cargo marked by ubiquitin for degradation is packaged into intraluminal vesicles with the help of the ESCRT complex, while the remaining cargo is recycled via different pathways to the plasma membrane or the Golgi apparatus. During these processes early endosomes mature into late endosomes^3–6^. Late endosomes then fuse with lysosomes to form endolysosomes, which are acidic and hydrolytically active, and serve as major sites for the degradation of cargo delivered by late endosomes^7^. Endolysosomes are characterized by the presence of RAB7 and LAMP1 on the membrane, along with an acidic lumen containing active acid hydrolases like cathepsins. After degradation of their content, endolysosomes need to undergo a maturation event to generate lysosomes again, which would be able to fuse with late endosomes to form new endolysosomes. This lysosome recycling is called *E*ndocytic *L*ysosome *R*eformation (ELR)^7–10^. During ELR, a tube emerges from the endolysosome, which undergoes fission and then becomes a lysosome. While there are reports on the tubule formation and fission in ELR, the underlying mechanisms especially the identity of the fission machinery remains elusive. Previous studies have implicated phosphoinositide signaling in ELR initiation. PIKfyve-generated PI(3,5)P_2_ on micro-domains of endolysosome drives tubulation^10^. Moreover, PI4KIIIβ delivered by the small GTPase ARF1 to the fission site provides PI4P necessary for the fission^9,10^. In contrast to ELR, Autophagic Lysosome Reformation (ALR) is a much better characterized lysosome reformation pathway. It depends on mTOR reactivation during prolonged starvation, initiating tubulation from autolysosomes^11^. PtdIns(4,5)P_2_ mediated KIF5B recruitment leads to the tubulation of the autolysosome^12^. WHAMM (WASP homolog associated with actin, Golgi membranes and microtubules), a member of the Wiskott-Aldrich syndrome protein (WASP) family driven Arp2/3-dependent actin remodeling facilitates the formation of tubules^13^. At last, PtdIns(4,5)P_2_ mediates the recruitment of clathrin and dynamin 2 to cause scission^14^. However, whether ELR uses the same cellular machinery as ALR remains unanswered.

In this study, we establish that ELR, unlike ALR, proceeds independently of both mTOR and dynamin 2 (DNM2). Unexpectedly, DRP1, a GTPase classically known for its role in mitochondrial and peroxisomal fission^15–17^, is essential for endolysosomal tubule fission. Fission sites are marked by both ER and mitochondria. Cells deficient in DRP1 or expressing a GTPase-dead mutant (DRP1-K38A) exhibit abnormally elongated endolysosomal tubules, indicative of impaired fission. A similar phenotype is observed when ER contact is disrupted. Surprisingly disruption of mitochondrial function leads to inhibition of the initiation of ELR, with no detectable tubulation. Using mitochondria-targeted calcium probes and specific inhibitors for endolysosomal and mitochondrial calcium channels, we demonstrate that calcium efflux from the reforming endolysosome to mitochondria is essential for the initiation of tubulation. We establish an order of events during ELR: First, initiation of ELR by Ca^2+^ flux from the endolysosome into mitochondria. Second, tubule formation and third scission of the lysosomal tubule through a concerted action of a tripartite membrane contact site of ER, mitochondria and the endolysosomal tubule, and the GTPase DRP1.

## Results

### Lysosomes reform from endolysosomes through three phenotypically distinct ways

In order to investigate ELR, we used an assay established by Bissig et al. (2017) to reversibly block ELR by the PIKfyve inhibitor YM201636. In brief, HeLa cells were treated with YM201636 for 2 hours to inhibit lysosome reformation. The drug was washed out, and cells were allowed to recover in drug-free media to synchronize the reformation events (Fig. 1A, Movie S1). During the recovery phase, the enlarged endolysosomes underwent reformation, leading to an increase in the number of LAMP1-positive structures over a 3 hr period. This increase was paralleled by an increase in lysotracker-positive structures (Fig. 1B). Quantitative analysis revealed that most reformation events occurred within the first hour post-washout for LAMP1-positive structures, while the number of lysotracker-positive structures continued to increase up to 2 hours, suggesting that newly formed proto-lysosomes were gradually acidified (Fig. 1C, D). To examine the reformation process in greater detail, we employed super-resolution live-cell imaging, capturing events at shorter time intervals. We observed three phenotypically distinct ways of lysosome reformation from endolysosomes: tubulation, budding, and shape distortion (Fig. 1E). Quantification of these events indicated that tubulation was the most prevalent mode, with nearly half of the reforming endolysosomes exhibiting tubules, followed by budding and shape distortion (Fig. 1F). To confirm the presence of three phenotypically different ways of ELR, we used two other methods to enlarge the endolysosome and followed the lysosome reformation. First, we enlarged the endocytic compartment using nigericin for 20 minutes followed by 20 minutes recovery in drug free medium^18^. Second, we employed the formation of sucrosomes by feeding sucrose to the cells overnight followed by the addition of invertase to cells to initiate lysosome reformation^7^. Under both conditions we could visualize the same three events i.e. tubulation, budding and shape distortion during ELR (Fig. S1A, B).

**Figure 1.**
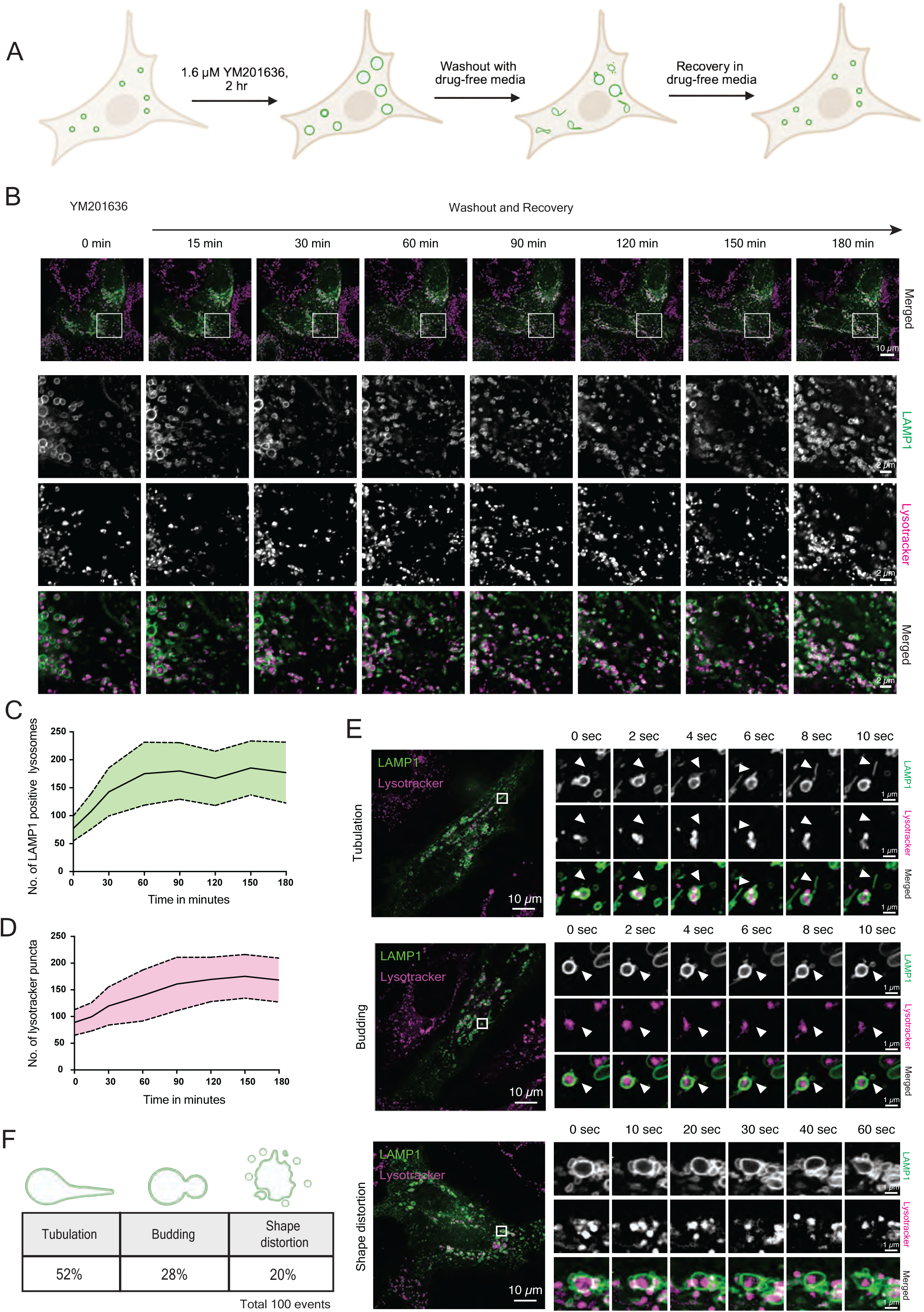
Live-cell imaging reveals the dynamics of endolysosomal reformation. (A) Schematic representation of the YM201636 treatment and washout used for synchronization of endocytic lysosomal reformation (ELR) events. (B) Live-cell imaging of cells expressing LAMP1-GFP (green) and stained with LysoTracker Deep Red (magenta), following washout of YM201636 and recovery in drug-free media for 3 h. Images were acquired every 15 min to monitor the timeline and progression of ELR events. (C) Quantification of LAMP1-positive structures from (B). The line plot depicts the number of LAMP1-positive compartments per cell over the recovery period. Data represent measurements from 30 cells. The central line indicates the mean; shaded regions or lines above and below represent the standard deviation (SD). (D) Quantification of LysoTracker-positive structures from (B). The line plot shows the number of LysoTracker-positive compartments per cell over the recovery period. Data represent measurements from 30 cells. The central line indicates the mean; shaded regions or lines above and below represent the SD. (E) Super-resolution imaging of endolysosomes undergoing reformation using spinning disk confocal microscopy equipped with SoRa Disk. Cells expressing LAMP1-GFP (green) were treated with YM201636 for 2 h and imaged immediately after washout in drug-free media to capture ELR events. Images were acquired over 2 min at 2 sec intervals. White arrowheads indicate sites of fission and ELR events. (F) Quantification of data from (E). Table summarizing the percentage occurence of three distinct types of reformation events observed out of the first 100 events from 30 cells.

It has previously been shown that PIKfyve inhibition and subsequent lysosome reformation after washout does not elevate autophagy in the cells^10,19^. To ensure that we were not inducing ALR in our assay, we assessed the accumulation LC3 by immunofluorescence in cells after YM201636 treatment and washout. Nutrient starvation was used as a positive control. We only observed the LC3 accumulation in starved cells but not in our PIKfyve inhibition conditions (Fig. S1C). Likewise, autophagic flux was not significantly increased (Fig. S1D). Together these results suggest that endolysosomes reform into lysosomes predominantly through tubulation, and confirm that YM201636 treatment and washout serves as a reliable tool to study these events.

### ELR does not require de novo protein synthesis

We observed an increase in the number of lysosomes following the washout of YM201636. To ensure that this increase was not due to de novo lysosomal biogenesis, we examined the levels of the lysosomal membrane protein LAMP1 and the luminal hydrolase cathepsin D. No significant changes in the levels of LAMP1 or cathepsin D were detected between untreated cells, YM201636 treated cells and those allowed to recover in drug-free medium for 1 hr (Fig. 2A, B). To further exclude the possibility of de novo lysosome formation during recovery, we added the translation inhibitor cycloheximide in the recovery medium. Live-cell imaging of endolysosomal reformation (ELR) revealed that cells recovering in the presence of cycloheximide exhibited similar lysosomal reformation dynamics to those in vehicle-treated controls (Fig. S2A, B). These results suggest that the observed increase in lysosome number may arise from recycling of pre-existing lysosomal membrane proteins and hydrolases rather than synthesis of new proteins.

**Figure 2.**
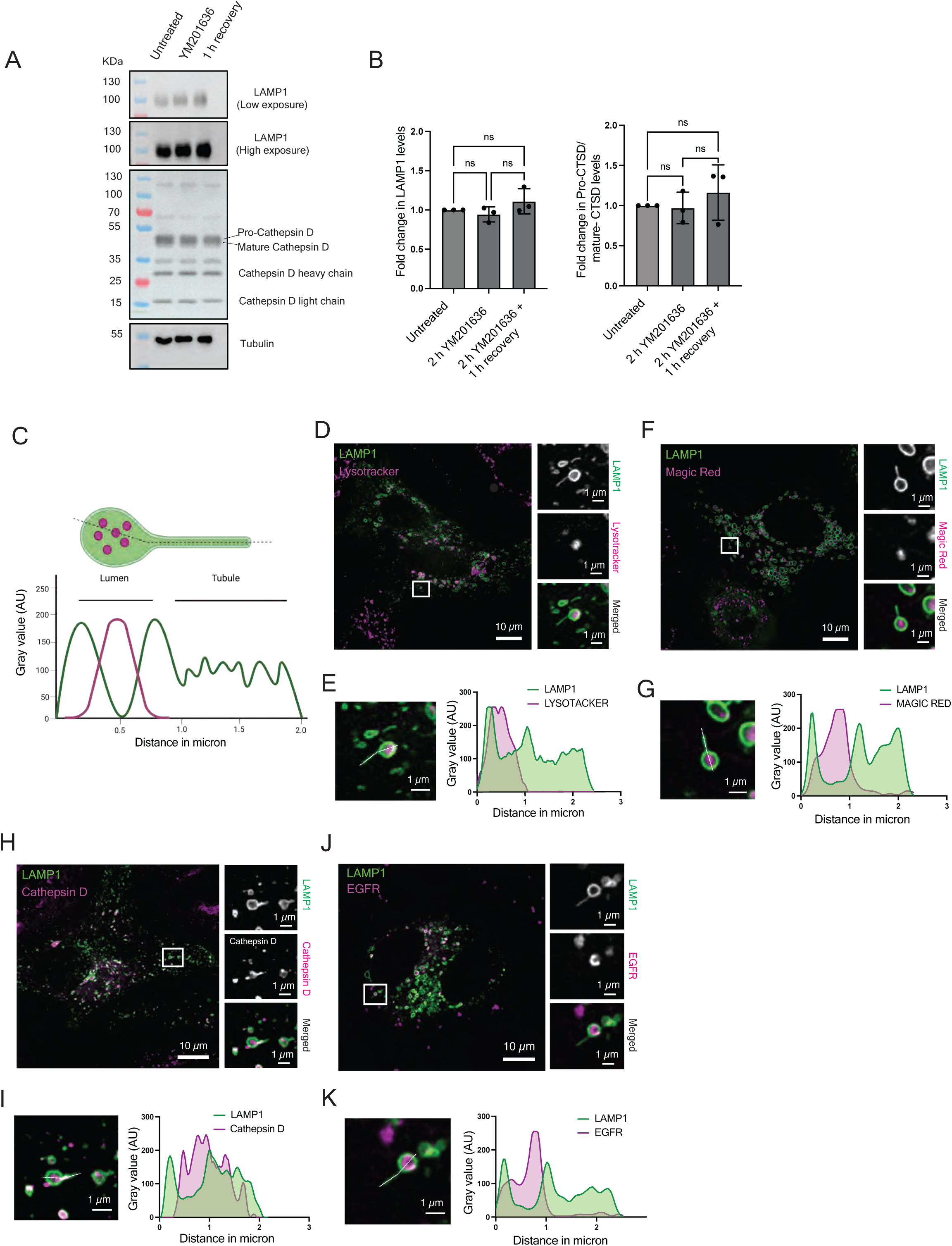
Tubulating lysosomes enable cargo sorting during reformation. (A) Immunoblot analysis of LAMP1 and Cathepsin D during reformation, using tubulin as a loading control. (B) Quantification of (A). Protein levels of LAMP1 and Cathepsin D at different maturation stages were analyzed for untreated, during YM201636 treatment, and post 1 h recovery. Fold change was determined by comparison to untreated control. Data represent mean of n=3 replicates; one-way ANOVA using Tukey’s multiple comparison, ns P = 0.8081 (Untreated vs 2 h YM201636), ns P= 0.4754 (Untreated vs 1 h recovery), ns P=0.2265 (2 h YM201636 vs 1 h recovery). For CTSD, ns P= 0.9871 (Untreated vs 2 h YM201636), ns P= 0.6745 (Untreated vs 1 h recovery), ns P=0.5877 (2 h YM201636 vs 1 h recovery) (C) Schematic representation of the analysis for cargo transport in tubules. (D) Super-resolution imaging of a tubulating endolysosome from a cell expressing LAMP1-GFP (green) and stained with Lysotracker Deep Red (magenta) using a spinning disk confocal with SoRa disk. Cells were treated with YM201636 for 2 h followed by washout and recovery to initiate the ELR. (E) Intensity plot for (D), showing intensity values of green and magenta channels over distance (μm). (F) Super-resolution image of a tubulating endolysosome from a cell expressing LAMP1-GFP (green) and stained with Magic Red (magenta) using a spinning disk confocal with SoRa disk. Cells were treated with YM201636 for 2 h followed by washout and recovery to initiate the ELR. (G) Intensity plot for (F), showing intensity values of green and magenta channels over distance (μm). (H) Super-resolution image of a tubulating endolysosome from a cell expressing LAMP1-GFP (green) and Cathepsin D-RFP (magenta) using a spinning disk confocal with SoRa disk. Cells were treated with YM201636 for 2 h followed by washout and recovery to initiate the ELR. (I) Intensity plot for (H), showing intensity values of green and magenta channels over distance (μm). (J) Super-resolution image of a tubulating endolysosome from a cell expressing LAMP1-GFP (green) and EGFR-mCherry (magenta) using a spinning disk confocal with SoRa disk. Cells were treated with YM201636 for 2 h followed by washout and recovery to initiate the ELR. Cells were also stimulated with EGF for 30 min to increase EGFR uptake. (K) Intensity plot for (J), showing intensity values of green and magenta channels over distance (μm).

### Endolysosomal tubules contain lysosomal proteins but are not acidified

Next, we explored whether we could monitor lysosomal activity and recycling of lysosomal proteins during ELR our system (Fig. 2C). When we stained cells with LysoTracker, a clear segregation of fluorescence was observed between the globular part and the tubules: LysoTracker accumulated in the globular region, whereas the tubules were devoid of signal (Fig. 2D, E). Similarly, hydrolase activity was confined to the globular part as determined by the protease activity marker Magic Red (Fig. 2F, G). Live-cell imaging of tubule fission further confirmed the absence of LysoTracker and Magic Red in the newly formed proto-lysosomes (Fig. S2C, D), indicating that these structures are not acidified and hence lack hydrolase activity. To visualize the redistribution of luminal cargo during ELR, we expressed RFP-tagged cathepsin D and followed its localization during tubulation and fission. Cathepsin D fluorescence was detectable in both the luminal and tubular regions of the tubulating endolysosome (Fig. 2H, I), and live imaging confirmed its presence in fissioned tubules (Fig. S2F, Movie S2). In contrast, when we tracked the fate of endocytic cargo, such as EGFR stimulated with EGF, the signal was restricted to the globular part and excluded from the tubules (Fig. 2J, K; Fig. S2E). Thus, while degradative activity remains confined to the endolysosomal globular part, lysosomal membrane proteins and hydrolases are transferred into degradation inactive tubules, which ultimately give rise to nascent proto-lysosomes.

### ELR and ALR are mechanistically distinct processes

Autophagic lysosomal reformation (ALR) is a well characterized pathway responsible for lysosome regeneration from autolysosomes. This process is known to depend on mTOR activity, and its inhibition has been shown to abrogate autolysosomal tubulation and fission^11^. To determine whether mTOR activity is required for ELR, we examined the phosphorylation status of S6 and S6K, two well-established downstream effectors of mTOR signaling. Cells treated with the mTOR inhibitor torin for 2 hrs served as a control for mTOR inactivation. Western blot analysis revealed no significant changes in the levels of phospho-S6 or phospho-S6K during YM201636 treatment or throughout the recovery period (Fig. 3A, B), consistent with previous findings on sucrosomes^20^. To further confirm that ELR occurs independently of mTOR activity in our system, we kept torin in the medium during the recovery period. Quantitative analysis of LAMP1-positive compartments after 30 minutes of recovery showed no significant difference in size between cells recovering in torin-containing medium and those in vehicle control (Fig. 3C, D). Similarly, the number of LAMP1-positive structures increased to a comparable extent in both conditions (Fig. S3A, B). Together, these results indicate that, unlike autophagic lysosome reformation (ALR), ELR proceeds independently of mTOR activity.

**Figure 3.**
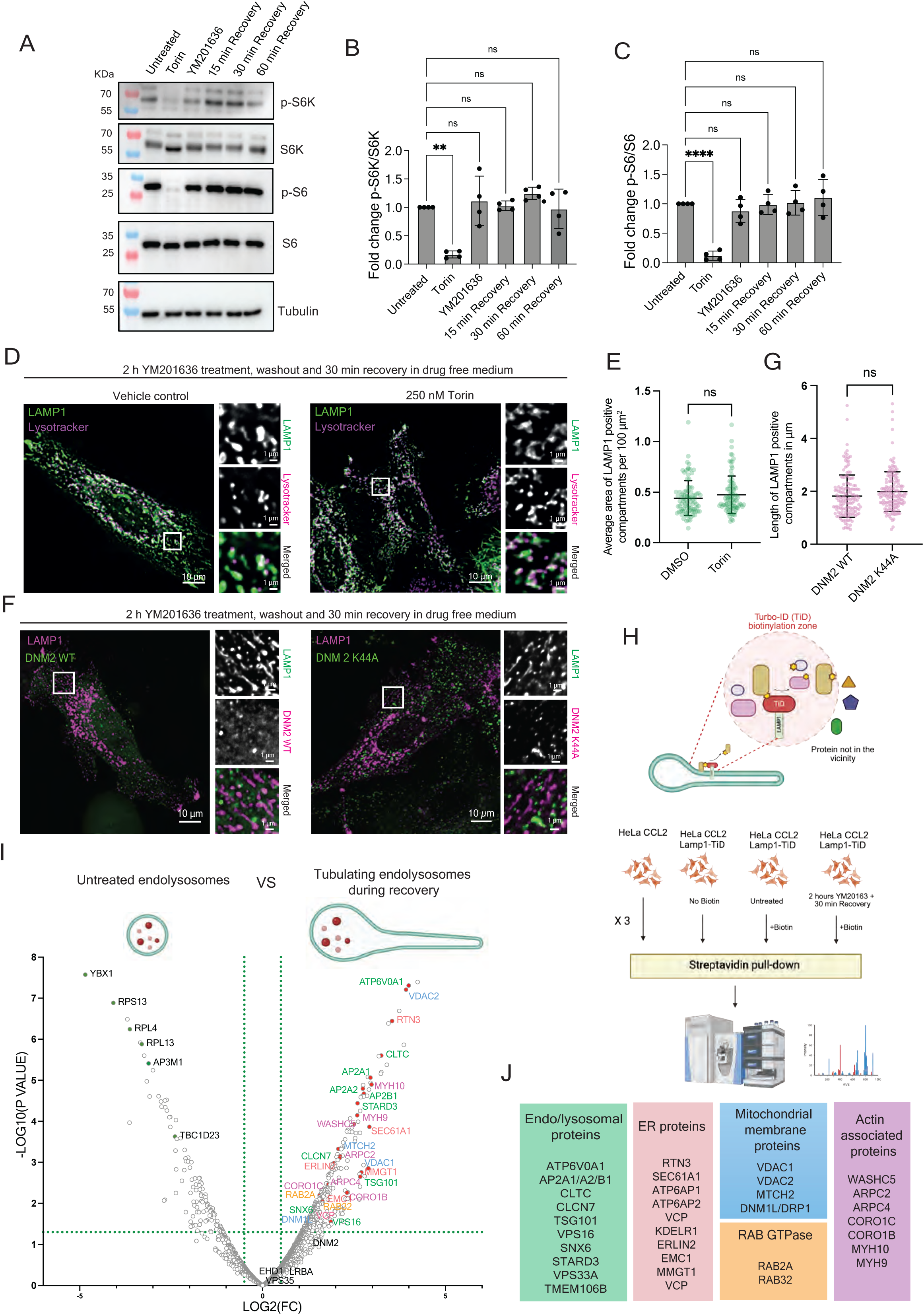
ELR progesses independently of mTOR activity and dynamin 2. (A) Immunoblot for the phosphorylation of mTOR effector proteins during different conditions. Torin was used as a positive control for mTOR inhibition. Tubulin was used as a loading control. (B) Quantification of the fold change p-S6K/S6K from immunoblot data in (A). Data represents the mean of n=4 replicates. one-way ANOVA using Tukey’s multiple comparison, **P= 0.0011 (Untreated VS Torin), ns P = 0.9809 (Untreated VS YM201636), ns P = 0.99997 (Untreated VS 15 min recovery), ns P = 0.6835 (Untreated VS 30 min recovery), ns P = 0.99997 (Untreated VS 60 min recovery). (C) Quantification of the fold change of p-S6K/S6K for from immunoblot data in (A). Data represents the mean of n=4 replicates. one-way ANOVA using Tukey’s multiple comparison, ***P = 0.000038 (Untreated VS Torin), ns P = 0.9370 (Untreated VS YM201636), ns P = 0.99999 (Untreated VS 15 min recovery), ns P = 0.999993 (Untreated VS 30 min recovery), ns P = 0.9605 (Untreated VS 60 min recovery). (D) Live-cell imaging of cells expressing LAMP1-GFP (green) stained with LysoTracker deep red (magenta) to assess the lysosomal reformation in presence of 250 nM torin. Images were acquired after 2 h YM201636 treatment and 30 min washout. (G) Average area of LAMP1-positive structures was quantified. Each dot represents the average area of per 100 µm^2^ ROI; plot shows mean area from a total of 90 ROI from 30 cells across n=3 biological replicates; Unpaired t-test; ns, P = 0.1980 (DMSO vs Torin). (F) Live-cell imaging of cells expressing LAMP1-mScarlet (magenta) together with DNM2-GFP or the GTPase mutant DNM2 (K44A)-GFP to assess DNM2 function in tubule fission. Images were acquired after 2 h YM201636 treatment and 30 min. (G) Quantification of LAMP1-positive tubule length (magenta) in cells overexpressing wild-type or mutant DNM2. Each dot represents the length of a single tubule; plot shows mean length from a total of 150 tubules across n=3 biological replicates; Unpaired t-test; ns P = 0.0564 (DNM2 WT vs. DNM2 K44A). (H) Schematic representation of the proximity labelling assay using LAMP1 fused to the biotin ligase TurboID. (I) Volcano plot comparing the biotinylated proxisome of LAMP1-TiD in cells treated with YM201636 for 2 h plus 30 min recovery versus untreated controls. Proteins significantly enriched during tubulation and fission are highlighted on the right side of the plot. Data from n=3 biological replicates. (J) Major categories of candidate proteins enriched during tubulation and fission.

Dynamin-2 (DNM2) has been shown to be essential for the fission of autolysosomal tubules during ALR and disruption of DNM2 function resulted in elongated tubules due to fission deficiency^14,21^. To investigate whether the fission machinery involved in ELR is identical to the one in ALR, we examined the role of DNM2 in the fission of endolysosomal tubules. We expressed GFP-tagged wild-type DNM2 and the GTPase-deficient mutant DNM2 K44A in cells expressing LAMP1-mScarlet. Surprisingly, we observed no significant difference in the length of LAMP1-positive tubules between cells expressing wild-type DNM2 and those expressing the K44A mutant (Fig. 3E, F). To further confirm this observation, we utilized DNM2 knockout (KO) cells and assessed their ability to undergo ELR. We detected no significant difference in the length of LAMP1-positive tubules between DNM2 KO and wild-type cells (Fig. S3C, D). Together, these findings indicate that DNM2, a key mediator of fission during ALR, is dispensable for ELR, suggesting that a distinct fission machinery operates in the endocytic lysosome reformation pathway.

### LAMP1 proxisome reveals DRP1 as a candidate involved in ELR

Given the mechanistic differences between ELR and ALR, we decided to use an unbiased approach to identify proteins in the proximity of the endolysosomes in untreated and after 30 min of recovery from YM201636 treatment using LAMP1-Turbo-ID (TiD) (Fig 3H). As expected, comparative proteomic analysis of cells undergoing ELR versus untreated cells revealed an enrichment of endolysosomal proteins during washout. Importantly, membrane-associated proteins from the ER such as the reticulin RTN3, the AAA-ATPase VCP and components of ER membrane protein complex (EMC) like EMC1, MMGT1 were also enriched. Surprisingly, levels of mitochondrial membrane proteins such as VDAC1, VDAC2 and MTCH2 were likewise increased in the recovery fraction (Fig. 3I, J). Intriguingly, the dynamin-related protein DRP1 (also known as DNM1L), which is essential for mitochondrial fission^22^ was also enriched. In contrast, DNM2 levels were not elevated in the same sample, consistent with our findings above, indicating that DNM2 is dispensable for ELR (Fig. 3I, J). Together, these results suggest that ER and mitochondrial membranes are in close proximity of endolysosomes during ELR and identify DRP1 as a possible candidate for tubule fission.

### DRP1 causes fission of endolysosomal tubules during ELR

To elucidate a potential role of DRP1 in endolysosomal tubule fission, we first checked whether DRP1 would localize to the endolysosomal tubule. To this end, we expressed mCherry-tagged DRP1 in cells expressing LAMP1-GFP. We observed a marked increase in DRP1 recruitment to endolysosomes during the recovery phase following drug washout (Fig. S4A, B). Time-lapse imaging of endolysosomes undergoing tubulation and fission revealed that DRP1 localized at fission sites (Fig. 4A, B, Movie S3).

**Figure 4.**
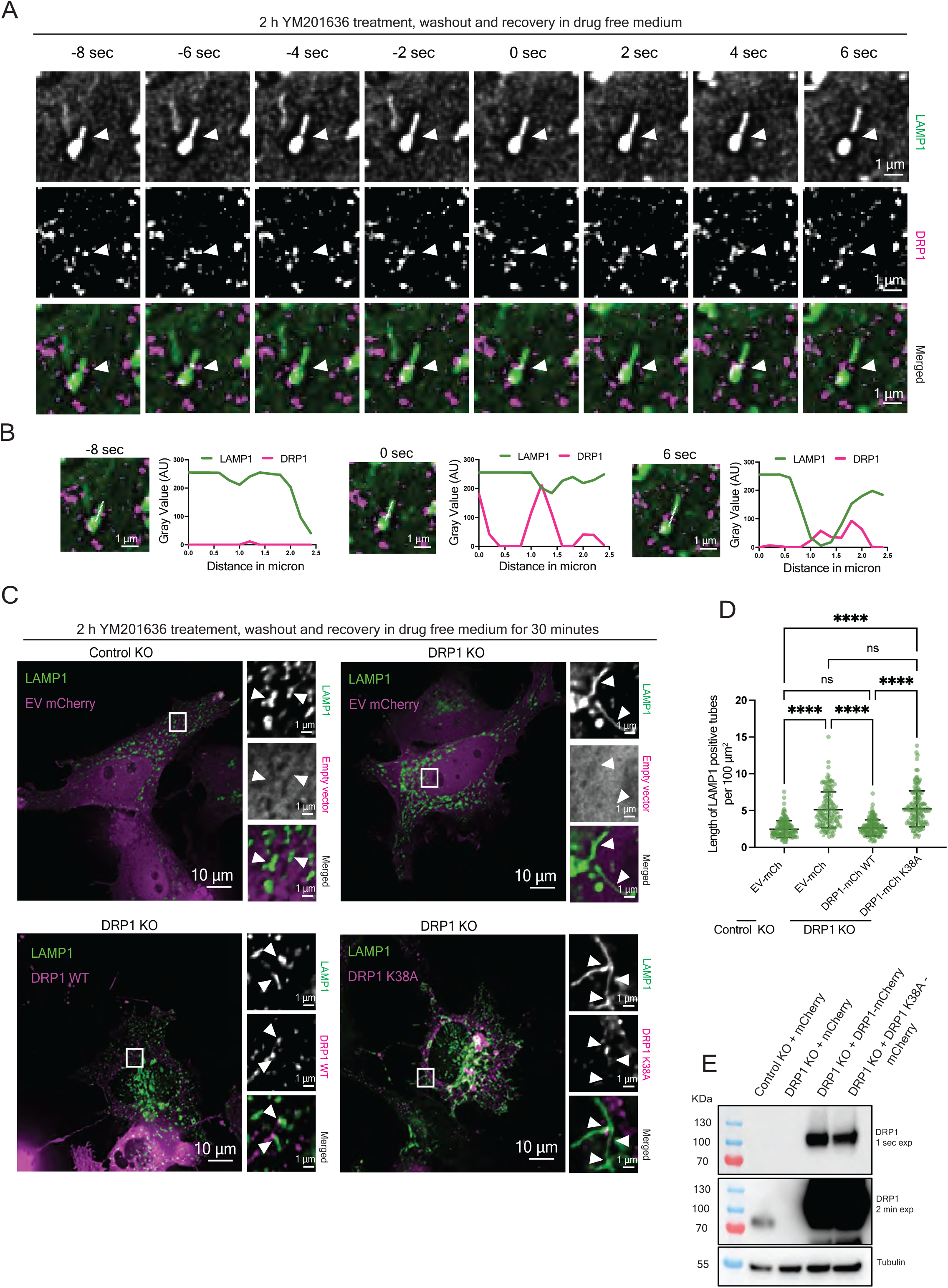
Endolysosomal tubule fission requires DRP1. (A) Live-cell confocal microscopy of cells expressing LAMP1-GFP (green) and DRP1-mCherry treated with YM201636 for 2 h, followed by washout and recovery in drug-free imaging buffer. Images were captured every 2 sec. White arrows indicate a LAMP1-positive tubule undergoing fission and DRP1 recruitment at the fission site. (B) Intensity plot for (A), showing green and magenta channel intensities across distance (μm) at three time points during fission events. (C) Imaging of LAMP1-positive tubules in wild-type and DRP1 knockout (KO) cells, as well as DRP1 KO cells overexpressing wild-type DRP1 or the DRP1 K38A mutant. Cells were treated with YM201636 for 2 h, followed by 30 min washout and recovery. White arrows indicate LAMP1-positive tubules and correspond to DRP1 localization. (D) Quantification of LAMP1-positive tubule length from (C). Each data point represents the length of a single tubule. Data from n=3 replicates; Kruskal-Wallis test with Dunn’s multiple comparisons: ****P < 0.1×10^−14^ (Control KO vs. DRP1 KO; DRP1 KO vs. DRP1 KO + WT DRP1; rescue with WT DRP1 vs. rescue with DRP1 K38A; Control KO vs. DRP1 KO + DRP1 K38A); ns, P > 0.9 ×10^−14^ (Control KO vs. DRP1 KO + WT DRP1; DRP1 KO vs. DRP1 KO + DRP1 K38A) (E) Immunoblot analysis confirming absence of DRP1 in DRP1 KO cells and overexpression of DRP1 in rescue experiments. Tubulin was used as a loading control.

To exclude the possibility that this recruitment was triggered by YM201636 treatment, we examined DRP1 localization in untreated cells and observed frequent colocalization of DRP1 with LAMP1-positive structures (Fig. S4C, D). To further assess the functional requirement of DRP1 in endolysosomal fission, we performed ELR assays in DRP1 knockout (KO) cells. During recovery, control HeLa cells displayed normal ELR with the formation of distinct proto-lysosomes, whereas DRP1 KO cells exhibited elongated LAMP1-positive tubules, indicating a defect in tubule scission. Re-expression of wild-type DRP1 in KO cells rescued this phenotype, while expression of the GTPase-deficient mutant DRP1 K38A failed to do so (Fig. 4C–E). Consistently, overexpression of DRP1 K38A in wild-type cells, even in the absence of YM201636 treatment, resulted in an accumulation of elongated LAMP1-positive tubules compared to wild-type DRP1 overexpression (Fig. S4E, F). As expected, similar results were obtained after induction of ELR (Fig. S4G, H, Movie S4). These results establish that DRP1 mediates the scission of endolysosomal tubules during lysosome reformation.

### Absence of DRP1 results in a reduction of lysosomal function

We wondered about the functional consequences lack of DRP1 on ELR and in particular on the lysosomal degradation capacity. To this end, we stained acidic compartments in control and DRP1 KO cells using LysoTracker Deep Red. DRP1 KO cells exhibited a reduction in both the total number and fluorescence intensity of LysoTracker-positive compartments when compared to the control (Fig. 5A, C). The distribution of LysoTracker-positive puncta was also markedly altered: in control, LysoTracker was present in two pools, a perinuclear and a peripheral pool distributed throughout the cell. In contrast, in DRP1 KO cells, LysoTracker signal was largely concentrated near the nucleus, and the peripheral pool was largely absent (Fig. 5B). To corroborate our observation, we next assessed lysosomal hydrolase activity using the cathepsin B substrate Magic Red. Consistent with the LysoTracker results, DRP1 KO cells displayed fewer Magic Red-positive compartments and reduced dye intensity relative to control cells (Fig. 5E, G). The distribution of Magic Red-positive puncta mirrored that of the LysoTracker pattern (Fig. 5F). Finally, DRP1 KO cells showed increased cholesterol accumulation, as detected by filipin staining, further reflecting impaired lysosomal function (Fig. 5H, J). Together, these results indicate that DRP1-dependent fission of endolysosomal tubules is crucial for maintaining a functional lysosomal pool within cells.

**Figure 5.**
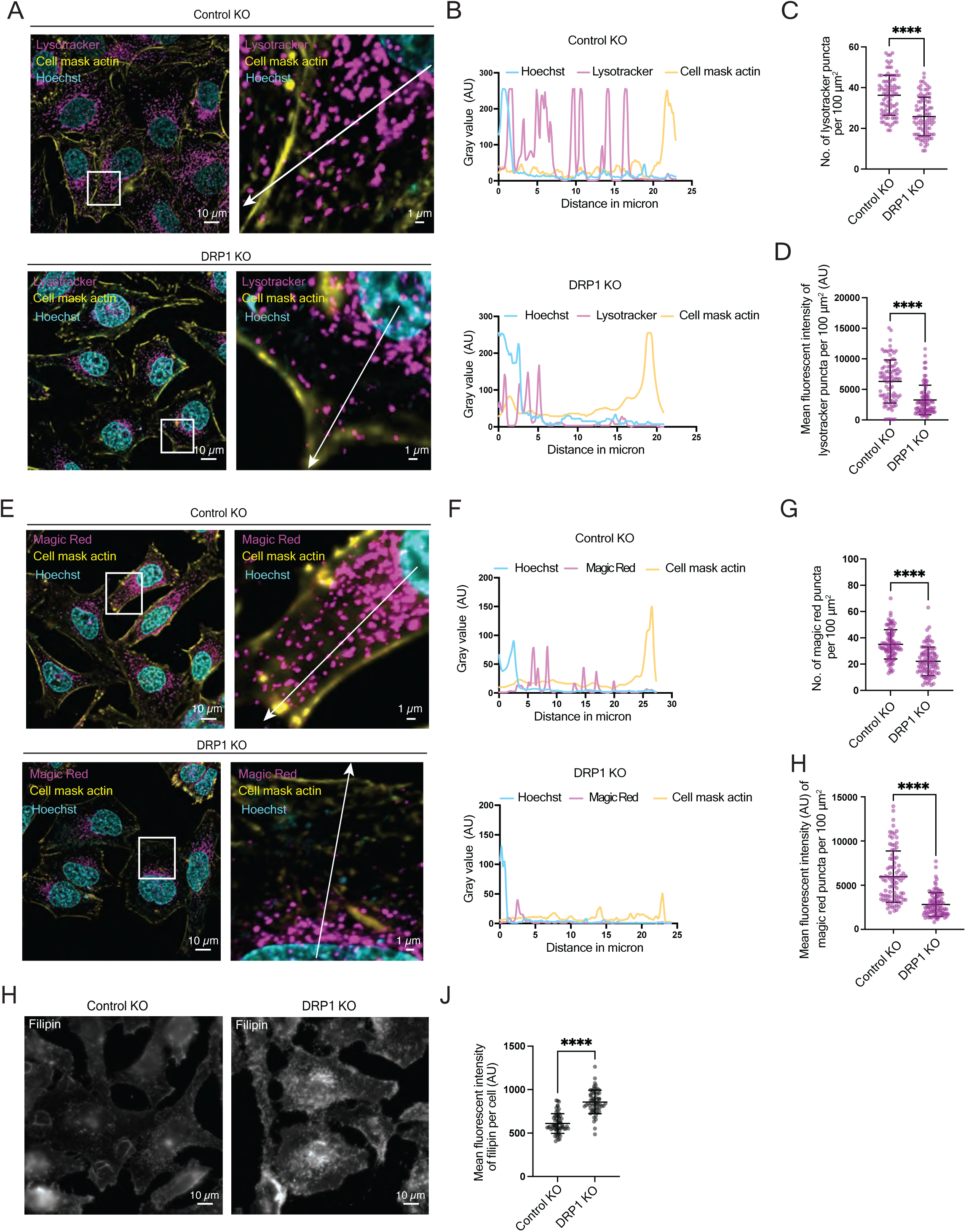
Loss of DRP1 impairs lysosomal function. (A) Live-cell imaging of LysoTracker Deep Red (magenta) in control knockout (KO) and DRP1 knockout (KO) cells to assess the effect of DRP1 KO on acidified compartments. Cells were also stained with Hoechst (cyan) to mark the nucleus and CellMask actin dye (yellow) to mark the cell periphery. The white arrow in the image indicates the region used for line plots. B) Intensity plot for (A), obtained by plotting intensities of Hoechst (cyan), LysoTracker Deep Red (magenta), and CellMask actin (yellow) across the distance in µm in the direction indicated by the arrowhead in (A). (C) Quantification of the number of LysoTracker puncta. Each dot represents the average number of LysoTracker-positive puncta (magenta) per 100 µm² ROI. The plot depicts the mean area from a total of 90 ROIs from 30 cells across n = 3 biological replicates. Unpaired *t*-test: ****P = 0.12096×10^−10^ (Control KO vs. DRP1 KO). (D) Quantification of the mean fluorescence intensity of LysoTracker puncta. Each dot represents the average intensity of LysoTracker-positive puncta (magenta) per 100 µm² ROI. The plot shows mean fluorescence intensity from a total of 90 ROIs from 30 cells across n = 3 biological replicates. Unpaired *t*-test: ****P = 0.304095×10^−9^ (Control KO vs. DRP1 KO). (E) Live-cell imaging of Magic Red (magenta) in control KO and DRP1 KO cells to assess the effect of DRP1 KO on cathepsin D activity in lysosomes. Cells were also stained with Hoechst (cyan) to mark the nucleus and CellMask actin dye (yellow) to mark the cell periphery. The white line in the image indicates the region used for line plots. (F) Intensity plot for (E), obtained by plotting intensities of Hoechst (cyan), Magic Red (magenta), and CellMask actin (yellow) across the distance in microns in the direction indicated by the arrowhead in (E). G) Quantification of the number of Magic Red puncta. Each dot represents the average number of Magic Red-positive puncta (magenta) per 100 µm² ROI. The plot shows the mean area from a total of 90 ROIs from 30 cells across *n* = 3 biological replicates; Unpaired *t*-test: ****P = 0.399×10^−12^ (Control KO vs. DRP1 KO). (H) Quantification of the mean fluorescence intensity of Magic Red puncta. Each dot represents the average intensity of Magic Red-positive puncta (magenta) per 100 µm² ROI. The plot shows mean fluorescence intensity from a total of 90 ROIs from 30 cells across n = 3 biological replicates; Unpaired *t*-test: ****P < 0.1×10^−14^ (Control KO vs. DRP1 KO). (I) Control KO and DRP1 KO cells were fixed and stained with filipin. The blue channel (filipin) is shown in the images. (J) Mean fluorescence intensity of the blue channel (filipin) was quantified and plotted as a dot plot. Each dot represents the gray value of one cell. The plot shows the mean fluorescence intensity of 30 cells from n = 3 biological replicates; Unpaired *t*-test: ****P < 0.1×10^−14^ (Control KO vs. DRP1 KO).

### ER and mitochondria mark the site for DRP1 recruitment on endolysosomal tubules

DRP1-dependent mitochondrial fission is preceded by an ER tubule contacting the fission site^23^. We detected ER and mitochondrial membrane proteins enriched during ELR in our LAMP1-TiD experiment (Fig. 3J). Therefore, we next examined whether DRP1 recruitment to endolysosomal tubules occurs at contact sites with the ER and/or mitochondria. We performed ELR assays in cells expressing LAMP1-GFP and DRP1-mCherry, and in which mitochondria were visualized using MitoTracker Deep Red or the ER by KDEL-BFP. DRP1 localized at the contact points between endolysosomal tubules and mitochondria (Fig. 6A, B), as well as between endolysosomal tubules and the ER (Fig. 6C, D), indicating the presence of the tripartite organellar contacts. To directly visualize the spatial organization of endolysosomal tubules with mitochondria and the ER, we used super-resolution microscopy and examined potential triple contact sites among these organelles. Strikingly, at sites where the ER contacted endolysosomal tubules, mitochondria were frequently present, suggesting the formation of tripartite contact sites during ELR (Fig. 6E, F). To further corroborate this observation and to determine whether these contacts occur specifically at fission sites, we performed live-cell confocal imaging of tubules undergoing scission. Both ER and mitochondria were observed in close apposition to endolysosomal tubules precisely at the fission site (Fig. 6G). Quantification of these events revealed that 59% of fission sites were simultaneously contacted by both ER and mitochondria. Interestingly, 29% of fission events occurred with only ER contact, 5% with mitochondria alone, and 7% without detectable association of either organelle (Fig. 6H). Together, these findings indicate that ER and mitochondria frequently cooperate at endolysosomal tubule fission sites, forming tripartite contact points that facilitate DRP1-mediated scission during lysosome reformation.

**Figure 6.**
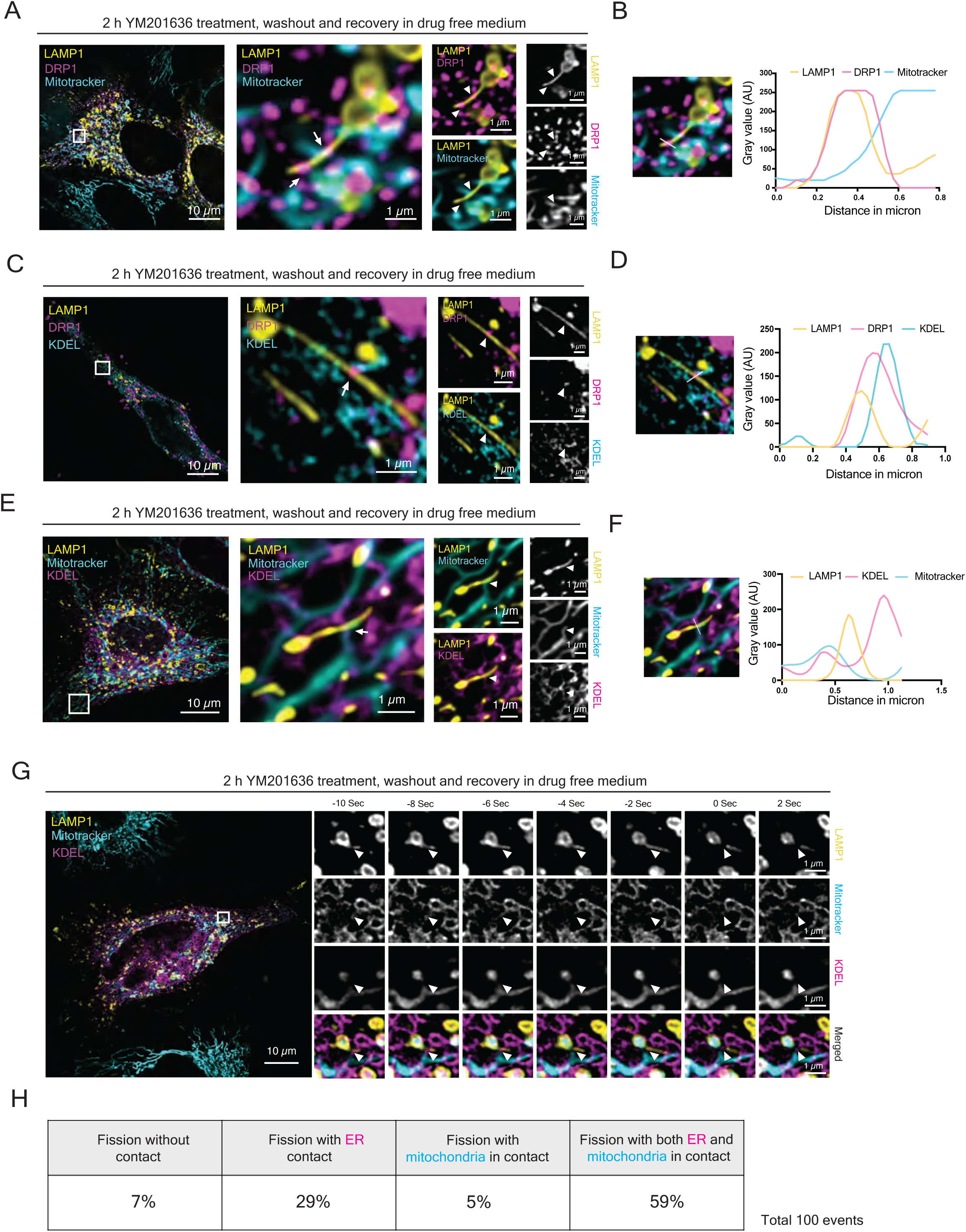
DRP1 recruitment to organelle contact sites co-ordinates endolysosomal fission. (A) Super-resolution imaging of endolysosomal tubules in cells expressing LAMP1-GFP (yellow), DRP1-mCherry (magenta), and stained with mitotracker deep red (cyan). Endolysosomal tubules were enriched using YM201636 and washout condition. (B) Intensity plot for data (A) obtained by plotting intensity values of LAMP1-GFP, DRP1-mCherry, and mitotracker channels over the distance in µm. (C) Super resolution imaging of endolysosomal tubules in cells expressing LAMP1-GFP (yellow), DRP1-mCherry (magenta), and KDEL-BFP (cyan). Endolysosomal tubules were enriched using YM201636 and washout condition. (D) Intensity plot for data (C) obtained by plotting intensity values of LAMP1-GFP, DRP1-mCherry, and KDEL-BFP over the distance in µm. (E) Super-resolution imaging of cells transfected with LAMP1-GFP (yellow), KDEL-BFP (magenta), and stained with mitotracker deep red (cyan). Endolysosomal tubules were enriched by YM201636 treatment for 2 h followed by washout and recovery in drug-free imaging buffer. White arrow indicates the contact site between ER, mitochondria, and endolysosomal tubule. (F) Intensity plots for data (E), showing intensities of all three channels along the line. Two intensity plots represent the proximity of three channels in two selected regions of interest (ROIs). (G) Live-cell super resolution microscopy shows contact sites exactly at the fission site. Cells were transfected with LAMP1-GFP (yellow), KDEL-BFP (magenta), and stained with mitotracker deep red (cyan). Fission events were enriched using YM201636 treatment for 2 h followed by washout and recovery. Cells were recorded for 2 min at 2 second intervals. The white arrowhead indicates the fission site. (H) Quantification of endolysosomal tubule fission events based on proximity to ER, mitochondria, both, or none. 100 fission events were counted from n=3 biological replicates.

### ER and mitochondrial morphology are required of ELR

The majority of fission sites of endolysosomal tubules involved ER and mitochondria. Next, we wanted to test whether these organellar contacts were functional relevant. Tubular ER is known to form contact with other membrane-bound organelles such as endosomes, mitochondria, lysosomes for their fission^9,23–25^. Reticulons play a role in maintaining tubular ER while CLIMP63 is involved in the sheet formation^26^. Overexpression of CLIMP63 has been implicated in increasing the sheet ER and reduction in tubules consequently reducing number of functional contact necessary for the fission^25,27^. Overexpression mCherry-tagged CLIMP63 in cells co-expressing LAMP1-GFP revealed significantly elongated LAMP1-positive tubules compared to KDEL-expressing controls after 30 min recovery from YM201636 treatment (Fig. S5A, B, F), suggesting that tubular ER contacting the endolysosomal tubule is a pre-requisite for efficient fission.

To perturb mitochondrial dynamics, we treated cells with the mitochondrial uncoupler CCCP, which induces hyper-fragmentation of mitochondria^28–30^. We performed an ELR assay in which during the recovery period 20 µM CCCP or DMSO as a control were present. Mitochondria were labeled with MitoTracker Deep Red. Strikingly, in CCCP-treated cells, endolysosomes even failed to initiate tubulation and remained enlarged, resembling the morphology observed during PIKfyve inhibition (Fig. S5C, D, G). These findings indicate that functional mitochondrial morphology and dynamics are essential for the initiation of endocytic lysosome reformation. Because CCCP treatment could also lead to energy depletion, we sought of a different way to perturb mitochondria and decided to impair the mitochondrial dynamics and function by knocking out the mitochondrial fission factor MFF. Oligomerization of MFF on the mitochondrial membrane is known to recruit DRP1 and cause fission of the mitochondria and hence the absence of MFF is known to cause hyper fusion of the mitochondria^28,31^. As expected, MFF-deficient cells displayed elongated, hyperfused mitochondria (Fig. 7A, S5E). During ELR, MFF KO cells retained enlarged endolysosomes resembling those in drug-treated conditions. Re-expression of MFF fully rescued this phenotype (Fig. 7A, C). Notably, endolysosomal enlargement was also evident in MFF KO cells even in the absence of YM201636 treatment (Fig. S5E, H), indicating a basal impairment in endolysosomal dynamics. Furthermore, expression of DRP1 in MFF-deficient cells prior to YM201636 treatment partially restored mitochondrial morphology and reduced endolysosomal enlargement after recovery (Fig. 7B, D). Taken together, our findings demonstrate that at least the ER contacts are essential for endocytic lysosome reformation (ELR). Perturbation of ER morphology and dynamics impaired endolysosomal tubule fission, underscoring the requirement of functional ER networks for efficient ELR. We hypothesize that mitochondria are similarly important in the process of tubule fission. Our data revealed, however, an additional function of mitochondria already in the initiation of ELR.

**Figure 7.**
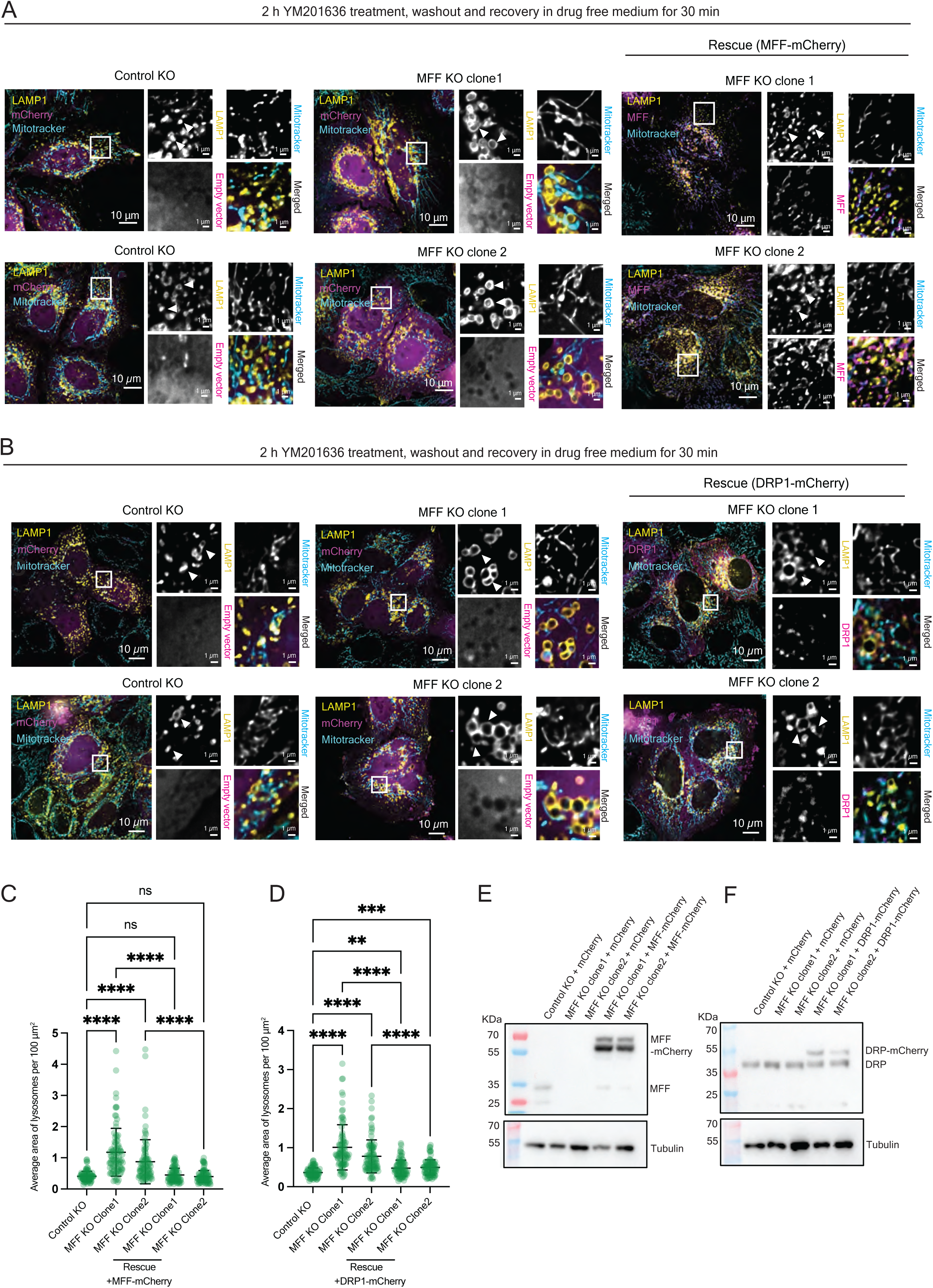
MFF KO effectuates inhibition of ELR. (A) Live-cell imaging of LAMP1-positive lysosomes (yellow) in control KO and MFF knockout (KO) cells, as well as MFF KO cells overexpressing MFF–mCherry (magenta). mCherry overexpression alone was used as a control for control KO and MFF KO conditions. Two different clones were imaged and shown. Cells were treated with YM201636 for 2 h, followed by 30 min washout and recovery. Cells were stained with MitoTracker Deep Red (cyan). White arrowheads highlight lysosomes in the respective conditions. (B) Live-cell imaging of LAMP1-positive lysosomes (yellow) in control KO and MFF KO cells, as well as MFF KO cells overexpressing DRP1 (magenta). mCherry overexpression alone was used as a control for control KO and MFF KO conditions. Two different clones were imaged and shown. Cells were stained with MitoTracker Deep Red (cyan) and treated with YM201636 for 2 h, followed by 30 min washout and recovery. White arrowheads highlight lysosomes in the respective conditions. (C) Quantification of the average area of LAMP1-positive structures for (A). Each dot represents the average area per 100 µm² ROI. The plot shows mean area from a total of 90 ROIs from 30 cells across n = 3 biological replicates. Kruskal-Wallis test with Dunn’s multiple comparisons: ****P < 0.1×10^−14^ (Control KO vs MFF KO clone 1), *****P = 0.323310×10^−9^ (Control KO vs MFF KO clone 2), ns P > 0.9 ×10^−14^ (Control KO vs MFF KO clone 1 rescue; Control KO vs MFF KO clone 2 rescue), ****P < 0.1×10^−14^ (MFF KO clone1 vs MFF KO clone 1 rescue), ****P = 0.2461 ×10^−11^ (MFF KO clone 2 vs MFF KO clone 2 rescue). (D) Quantification of the average area of LAMP1-positive structures for (B). Each dot represents the average area per 100 µm² ROI. The plot shows mean area from a total of 90 ROIs from 30 cells across n = 3 biological replicates; Kruskal-Wallis test with Dunn’s multiple comparisons: ****P < 0.1×10^−14^ (Control KO vs MFF KO clone 1; Control KO vs MFF KO clone 2), **P = 0.00847 (Control KO vs MFF KO clone 1 rescue), ***P = 0.00062 (Control KO vs MFF KO clone 2 rescue), ****P = 0.5×10^−14^ (MFF KO clone1 vs MFF KO clone 1 rescue), ****P=0.7456 ×10^4^ (MFF KO clone 2 vs MFF KO clone 2 rescue). (E) Immunoblot analysis confirming absence of MFF in MFF KO cells and overexpression of MFF in rescue experiments. Tubulin was used as a loading control. (F) Immunoblot analysis confirming absence of MFF in MFF KO cells and overexpression of DRP1 in rescue experiments. Tubulin was used as a loading control.

### Calcium efflux from lysosome to mitochondria initiates ELR

Therefore, we aimed to determine the nature of this ELR initiation function of mitochondria. Our proximity labeling data (Fig. 3J) indicated an enrichment of mitochondrial VDAC near endolysosomal tubules during ELR. We hypothesized that calcium signaling between endolysosomes and mitochondria might be required to initiate ELR. Calcium export from lysosomes through TRPML1 (Mucolipin-1) has previously been implicated in lysosomal fission^32^. TRPML1 is known to interact with VDAC (voltage-dependent anion channel) on the mitochondrial membrane to form functional contact sites that facilitate Ca^2+^ exchange between lysosomes and mitochondria^33^. To test the possibility that Ca^2+^ transfer from lysosomes to mitochondria might serve as a trigger for the initiation of ELR, we employed a mitochondrial Ca^2+^ sensor, mito-RCaMP1h and visualized changes in mitochondrial Ca^2+^ levels during ELR (Fig. 8A). Indeed, we observed a marked increase in the mito-RCaMP1h signal at 15 and 30 minutes after YM201636 washout, indicating a Ca^2+^ influx into mitochondria during the initiation of ELR (Fig. 8C, D). To establish a requirement of the TRPML1-VDAC axis in mediating this Ca^2+^ transfer, we used specific inhibitors of either TRPML1 or VDAC during ELR (Fig. 8B). Cells treated with the TRPML1 inhibitor ML-SI3, exhibited a pronounced defect in the initiation of ELR. In contrast, when ML-SI3 was only present during the YM201636 treatment, but absent during the recovery period, cells successfully initiated ELR. Interestingly, the inhibition was reversible. As soon as ML-SI3 was removed from the medium, ELR was initiated (Fig. 8E, F). Similarly, using the VDAC inhibitors Erastin or VBIT-4, resulted in persistent endolysosomal enlargement even after YM201636 removal. As expected, the effects of both Erastin and VBIT-4 were reversible upon washout and recovery in drug-free media (Fig. 8G–J). Together, these results highlight the crucial role of Ca^2+^ transfer from lysosomes to mitochondria via the TRPML1-VDAC axis in the initiation of ELR and suggest a dual role for mitochondria in ELR.

**Figure 8.**
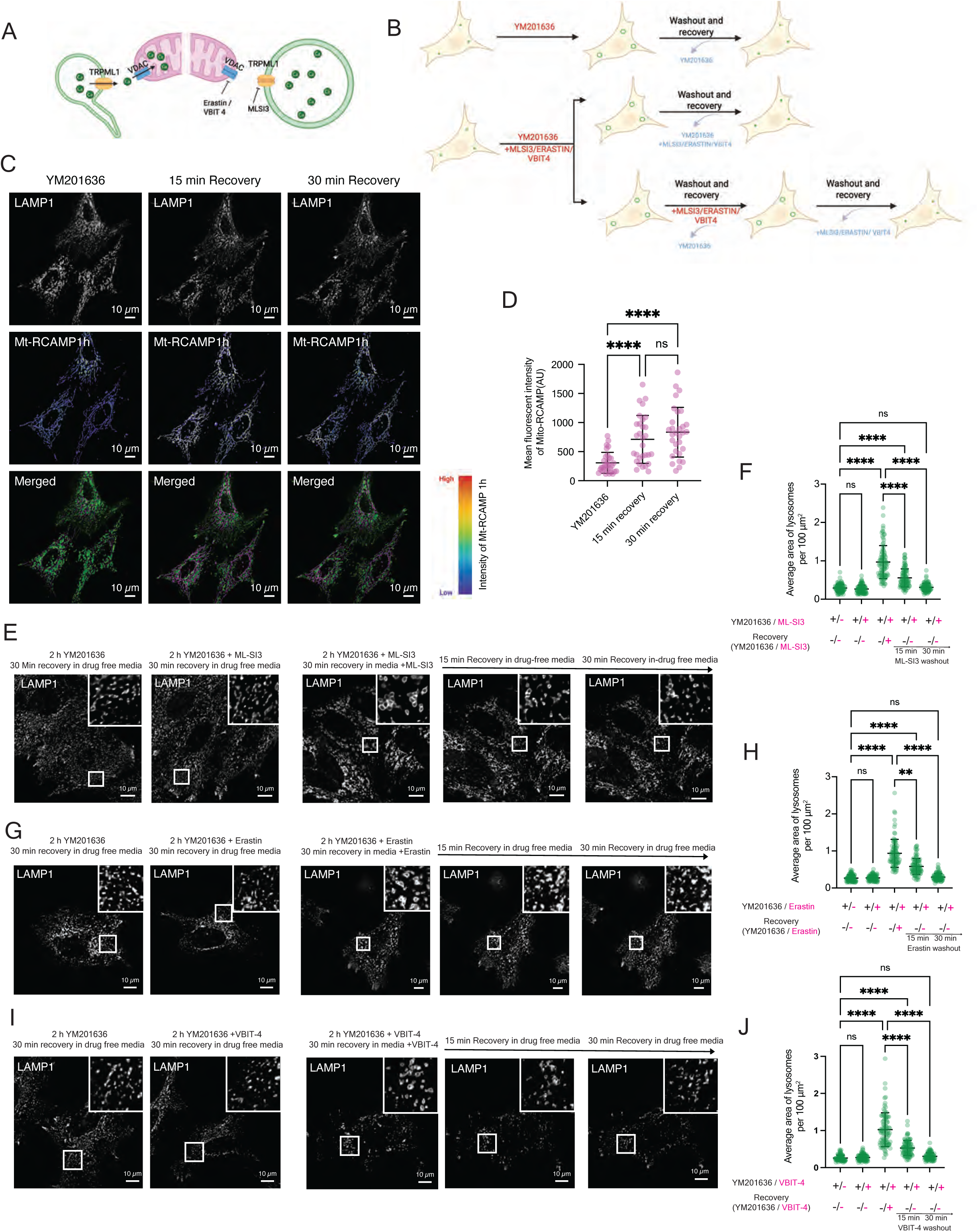
TRPML1–VDAC calcium transfer regulates lysosomal remodeling. (A) Schematic representation of calcium transport during ELR. (B) Schematic representation of the assay used to inhibit the lysosomal calcium exporter (TRPML1) or the mitochondrial calcium importer (VDAC) to assess its effect on ELR. (C) Live-cell imaging of cells co-transfected with LAMP1-GFP (green) and the mitochondrial calcium sensor mt-RCAMP1h (magenta). ELR was initiated with 2 h YM201636 treatment and washout. Cells were imaged for 30 min at 15-min intervals. D) Mean fluorescence intensity of mt-RCAMP1h (magenta) was quantified and plotted for 0 min, 15 min and 30 min recovery timepoints. Each dot represents the gray value of an individual cell. The plot shows the mean fluorescence intensity of 30 cells from n = 3 biological replicates; Kruskal-Wallis test with Dunn’s multiple comparisons: ****P = 0.5257 × 10^−4^ (YM201636 vs. 15 min recovery), ****P = 0.03522 × 10^−6^ (YM201636 vs. 30 min recovery), ns P > 0.9473 (15 min recovery vs. 30 min recovery). E) Live-cell imaging of cells expressing LAMP1-GFP and treated with YM201636 together with the TRPML1 inhibitor ML-SI3. Cells were subsequently washed and recovered in media containing only ML-SI3 to test the effect of ML-SI3 alone, as described in (B). ROIs show the LAMP1-positive compartments in representative parts of cells. (F) Quantification of the average area of LAMP1-positive structures for (E). Each dot represents the average area per 100 µm² ROI. The plot shows the mean area from a total of 90 ROIs from 30 cells across n = 3 biological replicates; Kruskal-Wallis test with Dunn’s multiple comparisons: ns P > 0.9 ×10^−14^ ( 2 h YM201636 + 30 min recovery vs. 2 h YM201636 and ML-SI3 + 30 min recovery), ****P < 0.1 ×10^−14^ (2 h YM201636 + 30 min recovery vs. 2 h YM201636 + 30 min recovery with ML-SI3), ****P = 0.691 × 10^−12^ (2 h YM201636 + 30 min recovery vs. 2 h YM201636 and ML-SI3 + 15 min recovery), ns P > 0.9 ×10^−14^ (2 h YM201636 + 30 min recovery vs. 2 h YM201636 and ML-SI3 + 30 min recovery), ****P = 0.4885 ×10^−4^ (2 h YM201636 + 30 min recovery with ML-SI3 vs. 2 h YM201636 and ML-SI3 + 15 min recovery), ****P = < 0.1 ×10^−14^ (2 h YM201636 + 30 min recovery with ML-SI3 vs. 2 h YM201636 and ML-SI3 + 30 min recovery). (G) Live-cell imaging of cells expressing LAMP1-GFP and treated with YM201636 together with the VDAC2/3 inhibitor erastin. Cells were subsequently washed and recovered in media containing only erastin to test the effect of erastin alone, as described in (B). ROIs show the LAMP1-positive compartments in representative parts of cells. (H) Quantification of the average area of LAMP1-positive structures for (G). Each dot represents the average area per 100 µm² ROI. The plot shows the mean area from a total of 90 ROIs from 30 cells across n = 3 biological replicates; Kruskal-Wallis test with Dunn’s multiple comparisons: ns P > 0.9 ×10^−14^ ( 2 h YM201636 + 30 min recovery vs. 2 h YM201636 and Erastin + 30 min recovery), ****P < 0.1 ×10^−14^ (2 h YM201636 + 30 min recovery vs. 2 h YM201636 + 30 min recovery with Erastin), ****P < 0.1 ×10^−14^ (2 h YM201636 + 30 min recovery vs. 2 h YM201636 and Erastin + 15 min recovery), ns P > 0.9 ×10^−14^ (2 h YM201636 + 30 min recovery vs. 2 h YM201636 and Erastin + 30 min recovery), **P = 0.00103 (2 h YM201636 + 30 min recovery with Erastin vs. 2 h YM201636 and Erastin + 15 min recovery), ****P = < 0.9 ×10^−14^ (2 h YM201636 + 30 min recovery with Erastin vs. 2 h YM201636 and Erastin + 30 min recovery). (I) Live-cell imaging of cells expressing LAMP1-GFP and treated with YM201636 together with the VDAC1 inhibitor VBIT-4. Cells were subsequently washed and recovered in media containing only VBIT-4 to test the effect of VBIT-4 alone, as described in (B). ROIs show the LAMP1-positive compartments in representative parts of cells. (J) Quantification of the average area of LAMP1-positive structures for (I). Each dot represents the average area per 100 µm² ROI. The plot shows the mean area from a total of 90 ROIs from 30 cells across n = 3 biological replicates; Kruskal-Wallis test with Dunn’s multiple comparisons: ns P > 0.9 ×10^−14^ ( 2 h YM201636 + 30 min recovery vs. 2 h YM201636 and VBIT-4 + 30 min recovery), ****P < 0.1 ×10^−14^ (2 h YM201636 + 30 min recovery vs. 2 h YM201636 + 30 min recovery with VBIT-4), ****P < 0.1 ×10^−14^ (2 h YM201636 + 30 min recovery vs. 2 h YM201636 and VBIT-4 + 15 min recovery), ns P =0.2364 (2 h YM201636 + 30 min recovery vs. 2 h YM201636 and VBIT-4 + 30 min recovery), **P = 0.29 ×10^−4^ (2 h YM201636 + 30 min recovery with VBIT-4 vs. 2 h YM201636 and VBIT-4 + 15 min recovery), ****P = < 0.1 ×10^−14^ (2 h YM201636 + 30 min recovery with VBIT-4 vs. 2 h YM201636 and VBIT-4 + 30 min recovery).

**Figure 9.**
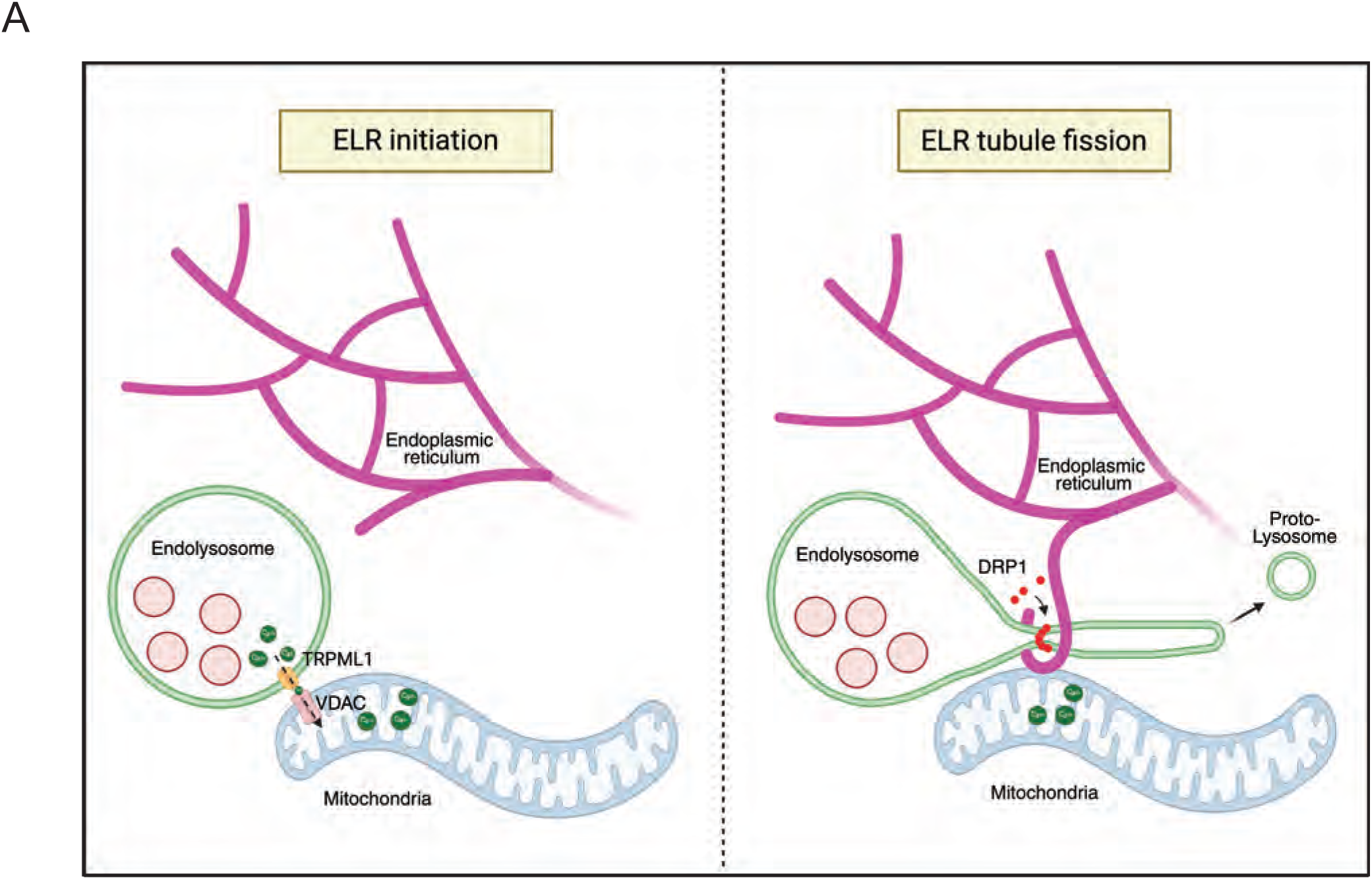
Schematic highlighting the role of ER and mitochondria contact in initiation and tubule fission of endolysosome during endocytic lysosome reformation. Endocytic lysosome reformation is initiated by calcium efflux from reforming endolysosomes to mitochondria at ER-endolysosome-mitochondria contact sites, which is required for tubule formation. DRP1 is recruited to these tripartite contact sites and localizes along endolysosomal tubules. At sites of ER contact, DRP1 mediates tubule scission to generate nascent lysosomes.

## Discussion

Endocytic lysosome reformation (ELR) has emerged as a critical process to maintain lysosomal homeostasis following the degradation of endocytic cargo. This process involves the generation of tubulo-vesicular compartments from acidified endolysosomes that recycle membranes and lysosomal membrane proteins into recycled proto-lysosomes^9,10,20^. Although the morphological stages of ELR have been described, the molecular machinery governing endolysosomal tubules formation and scission has remained unclear. Here, we delineate the underlying mechanism of tubular dynamics and identify key factors that regulate these events. Remarkably, even though ELR and ALR follow morphologically similar pathways with tubule formation and scission, the machinery required for ELR appears to be distinct from the one driving ALR. ALR is initiated by the reactivation of mTORC^11,34^, which has no detectable role in ELR. In contrast, ELR initiation requires Ca^2+^ release from endolysosomes and Ca^2+^ uptake by adjacent mitochondria. Moreover, while autolysosomal tubule fission to generate proto-lysosomes is dependent on dynamin 2^21^, ELR utilizes the dynamin-related protein DRP1. Finally, ELR requires contacts with the ER and mitochondria for the fission of endolysosomal tubules.

We assume that the mechanism of ELR and ALR are different because ELR is a continuous process that happens all the time and which is essential for cellular homeostasis, while ALR is part of a stress response. In support of this notion is the finding that in response of lysosomal damage with LLOMe, the ALR machinery is recruited and DNM2 is required for fission of lysosomal tubules during lysosomal membrane regeneration^35^. Therefore, the signals to initiate the reformation process are likely to be different. For ALR after starvation, reactivation of mTORC has been identified^11,34^. However, in lysosomal damage repair, mTORC1 is not required but rather LC3 and LC3-binding proteins are recruited, which may in turn mediate the recruitment of the ALR machinery for repair (Bhattacharya et al., 2023). During endocytic lysosome reformation, there is no damage and hence no repair or stress response is activated. Still a signal for initiation of the process must be provided. Lysosomal Ca²⁺ export via TRPML1 has been implicated in lysosome fission^32^, and mitochondria-lysosome contact through TRPML1-VDAC interaction in known to regulate this Ca^+2^ dynamics^33^. We suggest that the TRPML1-VDAC mediated Ca^2+^ flux from endolysosomes to the mitochondria could act as such a signal, and that a contact between endolysosomes and mitochondria should exist at least during the initiation of ELR. This Ca^2+^ is likely to induce Ca^2+^ signaling leading to the activation and/or recruitment of DRP1 to these sites^36^. In support of such an activation mechanism, we find that KO of the mitochondrial DRP1 receptor MFF also blocks ELR. DRP1-mediated fission of mitochondria is dependent on ER tubules that locally constrict and perturb the lipid bilayer and thereby allow the oligomerization of DRP1 to drive scission^23,37^. During ELR, the ER tubule does not contact mitochondria but the endolysosomal tubule. The ER contact with endolysosomal tubule during fission has been previously describe for phospholipid transfer across the membranes^9^. What drives this selectivity remains unclear. There is also the possibility that DRP1 is recruited to the ER and transferred from there to the endolysosomal tubule. ER tubulation is itself facilitated by the GTPase independent activity of DRP1 associated with ER^38^. Further, DRP1 can oligomerize at ER and can be transferred to the mitochondria for the fission^39^. When assessed the localization of DRP1 with ER / Mitochondria on the endolysosomal tubules, DRP1 localized at sites where both ER and mitochondria contacted the tubulating endolysosome, suggesting that these organelles mark the fission site may be through locally concentrating DRP1 at the prospective fission site.

The proto-lysosomes should contain at least some lysosomal markers. The endolysosomal tubules contain LAMP1 and assembled V-ATPase ^7,20^. We also detected a soluble hydrolase, cathepsin D in the tubules. However, neither degradative cargo, such as EGFR, nor active cathepsin B, as indicated by Magic Red were found in the endolysosomal tubules. Likewise, the tubules were negative for LysoTracker indicating that they were not acidified. Therefore, the degradative capacity appears to spatially restricted to globular part of the endolysosome. Moreover, degradative cargo seems to be confined to the globular part while hydrolases may diffuse or be actively sorted into the tubular part. Our data indicate that hydrolases in the tubular part of the endolysosome might no longer be active, Our study elucidates key aspects of the endocytic lysosome reformation pathway: ELR occurs through a fundamentally mechanistically distinct process than ALR, despite the morphological resemblance. Initiation of ELR requires Ca^2+^ signaling between endolysosome and mitochondria at contact sites that contain TRPML1 and VDAC. ELR is dependent on a tripartite organellar contact between ER, mitochondria and endolysosomal tubules and requires DRP1-mediated fission. Nevertheless, several questions remain unresolved. For example, the contributions of the cytoskeleton to endolysosomal membrane remodeling are not established. Such studies may provide insight into how tubulation and fission are coordinated. Moreover, the fate and maturation of proto-lysosomes into functional lysosomes still remain enigmatic.

## Methods

### Cell culture

Hela CCL2 cells (Gift from Dr. Martin Spiess) control, DRP1 knockout and DNM2 knockout HeLa cells (Gift from Dr. Mike Ryan) were grown in in high-glucose Dulbecco’s modified Eagle’s medium (Sigma-Aldrich) with 10% foetal bovine serum (FBS, Biowest), 2 mM L-glutamine, 100 U per mL penicillin G and 100 ng per mL streptomycin, 1 mM sodium pyruvate at 37 °C and 5% CO2. For starvation experiments cells were grown in HBSS (Gibco). All the cell lines used in this study were routinely tested for mycoplasma. For transient cell transfections, cells were plated into six-well plates to reach 70% confluency the following day and transfected with 1 µg plasmid DNA complexed with Helix-IN transfection reagent (OZ Biosciences).

### Generation of MFF-KO CRISPR/CAS9 cell line

HeLa CCL2 cells deficient of MFF were generated using the CRISPR/CAS9 system. gRNA as described previously by Pangou et al (2021)^40^ were used-guide 1-TAGTCGAATTCAGTACGAAA, guide 2-TGATAATGCAAGTTCCGGAG. Guides were cloned into pX458 (Addgene #48138) and pX459 (Addgene #62988) using the BbsI cloning site. Briefly, 1 × 10^6^ HeLa cells were seeded per 10-cm dish. The following day, cells were transfected with 6 μg DNA (3 μg/plasmid) with Helix IN transfection reagent (#HX10100; OZ Biosciences). For control cells, control vectors without gRNA insert were transfected. Puromycin resistant cells were selected with 1.5 mg/mL puromycin for 18 h. FACS sorting was carried out 48 h after transfection. Cells were trypsinized and resuspended in cell-sorting medium (2% FCS and 2.5 mM EDTA in PBS) and sorted on a BD FACS Aria Fusion Cell Sorter. GFP+ cells were collected and seeded into a 96-well plate. Cells were then expanded and MFF expression was determined by western blot analysis.

### Reagents and antibodies

The following reagents were used in this study: YM-201636 (Cat. HY-13228/ MCE), Bafilomycin (Cat. BML-CM110-0100/ enzo live sciences), Halt phosphatase inhibitor (Cat. 78420/ Thermo Fisher Scientific), Biotin (Cat. B450/ Sigma), Cycloheximide (Cat. C1988-1G/ Sigma), LysotrackerTM Deep Red (Cat. L12492/ ThermoFisher), Magic Red™ Cathepsin B Kit (Cat. ICT938/ Bio-Rad), MitoTracker Deep Red FM (Cat. M22426/ ThermoFisher), Filipin complex ready-made solution (Cat. SAE0088/ Sigma), CellMask™ Green Actin Tracking Stain (Cat. A57243/ ThermoFisher), Hoechst 33342 Fluorescent Nucleic Acid Stain (Cat# 639/ ImmunoChemistry technologies). Erastin (Cat. HY-15763/ MCE), VBIT-4 (Cat. HY-129122/MCE), ML-SI3 (Cat. HY-139426/ MCE), CCCP (Cat. C2759/ Sigma), Nigericin (Cas. 28643-80-3/ Adipogen), Invertase (I450-1G/ Sigma), Torin (Cat. HY-130002/ MCE).

The following antibodies were used in this study: Anti-LAMP1 (C54H11) Rabbit mAb(Cat. 3243S/ Cell Signaling Technology), Anti-Cathepsin D antibody (EPR3057Y) (Cat. ab75852/ Abcam, Anti α-Tubulin mouse mAb (Cat. T5168/ Sigma-Aldrich), Anti-P70 S6 Kinase (49D7) Rabbit mAb (Cat. 2708S/ Cell Signaling Technology), Anti-P-p70 S6 Kinase (T389) (108D2) Rabbit mAb (Cat. 9234L/ Cell Signaling Technology), Anti-S6 Ribosomal Protein (5G10) Rabbit mAb (Cat. 2217L/ Cell Signaling Technology), Anti-P-S6 Ribosomal Protein (S240/244) (D68F8) Rabbit mAb (Cat. 5364S/ Cell Signaling Technology), Anti-DRP1 (C-terminal) Mouse Polyclonal antibody (Cat. 12957-1-AP/ Proteintech), Anti-MFF Mouse Polyclonal antibody (Cat. 17090-1-1AP/ Proteintech), Anti-LC3B (D11) XP® Rabbit mAb (Cat. 3868S/ Cell Signaling Technology), goat anti-mouse IgG, (H+L), HRP-coupled (Cat. 31430/Pierce/Invitrogen), goat anti-rabbit IgG, (H+L), HRP-coupled (Cat. 31460/ Pierce/Invitrogen), goat anti-rabbit IgG(H+L) secondary antibody, AlexaFluor488 (Cat. A-11034/ Invitrogen). All primary antibodies were used at the dilution of 1:1,000 for immunoblot and at a dilution of 1:200 for immunofluorescence. The following secondary antibody was used at a dilution of 1:10,000 for immunoblot and at a dilution of 1:500 for immunofluorescence.

### DNA and Plasmid sources

The following commercially available plasmids were obtained: LAMP1-GFP (Cat. 34831/ Addgene), LAMP1-mScarlet (Cat. 98827/ Addgene), pSpCas9(BB)–2A-GFP (pX458) (Cat. 48138/ Addgene), pSpCAS9 (BB) 2A-puro (pX459) (Cat. 48139/ Addgene), EGFR-GFP (Cat. 32751/ Addgene), WT Dynamin 2-GFP (Cat. 34686/ Addgene), GFP-Dynamin2 K44A (Cat. 22301/ Addgene), BFP-KDEL (Cat. 49150/ Addgene), mCh-Climp63 (Cat. 136293/ Addgene), pmCherry C1 MFF (Cat. 157760/ Addgene), pCAG-mito-RCaMP1h (Cat. 105013/ Addgene). EGFR-mCherry was cloned by replacing the GFP from EGFR-GFP (Cat. 32751/ Addgene) with mCherry. mCherry was amplified by PCR from the mCherry-Rab11 (Cat. 55124/ Addgene) plasmid and inserted in the EGFR-GFP backbone with NEBuilder HiFi Assembly cloning kit (#E5520S; New England Biolabs). The vector encoding 3×HA-TurboID (Addgene #107171) was used as a template to amplify the TurboID gene. LAMP1-TurboID was generated by replacing GFP in the LAMP1-GFP vector (Addgene #34831) with TurboID using the NEBuilder HiFi DNA Assembly Cloning Kit (NEB, #E5520S). pcDNA 3.1 mCherry was a kind gift from Dr. Martin Spiess. Cathepsin D-RFP was a kind gift from Dr. Ana Maria Lennon Dumenil. mCherry-DRP1 and mCherry-DRP1 K38A was a kind gift from Dr. Oishee Chakrabarty.

### Immunostaining

HeLa cells were seeded onto glass coverslips 24 h before fixation to allow proper attachment and growth. Cells were fixed 48 h after plating with 4% paraformaldehyde, permeabilized using 0.1% Triton X-100, and subsequently blocked in phosphate-buffered saline (PBS) supplemented with 5% fetal bovine serum (FBS) to prevent nonspecific binding. Following blocking, samples were incubated with the designated primary antibodies, then probed with Alexa Fluor–conjugated secondary antibodies. Finally, coverslips were mounted onto glass slides using either Fluoromount-G (Southern Biotech) or Vectashield and sealed with clear nail polish to preserve fluorescence.

### Microscopy

Fixed-cell coverslips were imaged on an inverted Axio Observer microscope (Zeiss) equipped with a Plan-Apochromat N 63×/1.40 oil DIC M27 objective, a Photometrics Prime 95B camera, and Zen 2.6 acquisition software. The respective filter cubes with standard specifications for EGFP were used to image Alexa Fluor 488.

For live-cell imaging, cells were seeded into 4-well imaging chamber slides (Ibidi) one day prior to imaging to achieve a confluency of approximately 70% at the time of acquisition. Immediately before imaging, the culture medium was replaced with a pre-warmed imaging buffer composed of phosphate-buffered saline (PBS) supplemented with 5 mM D-glucose, 1 mM CaCl_2_, 2.7 mM KCl, and 0.5 mM MgCl_2_. The buffer was further supplemented with 10% fetal calf serum (FCS) and penicillin-streptomycin to maintain cell viability and prevent contamination during imaging. Unless specified otherwise, cells were treated with 1.6 μM YM201636 in complete DMEM for two hours to inhibit lysosome reformation. Images were taken after drug washout and a 30-min recovery in pre-warmed imaging buffer. Images were acquired at 37°C with 5% CO_2_ using an inverted Axio Observer microscope (Zeiss) with a Plan-Apochromat N 63×/1.40 oil DIC M27 objective, a Photometrics Prime 95B camera, and Zen 2.6 acquisition software. Filters with standard specifications for EGFP, DsRed, and Cy5 were used for GFP, mCherry, and deep-red dyes, respectively.

For confocal spinning disk super-resolution microscopy, we employed an Olympus SpinSR (CSU-W1) with a 60×/1.42 NA oil-immersion objective using the SoRa disk. A 3.2× intermediate magnification (192× combined) was used to image lysosomal tubules in stills as well as in time-lapse mode. Z-stacks were acquired for single time points, whereas time-lapse imaging was performed at a single z-plane at an interval of two seconds. Cells were imaged in imaging buffer at 37°C with 5% CO_2_. Lasers with wavelengths of 405 nm, 488 nm, 561 nm, and 640 nm were used to illuminate BFP, GFP, mCherry/mScarlet, and deep-red dyes, respectively.

Images were deconvolved with Huygens Remote Manager using Good’s Roughness Maximum Likelihood Estimation, with 50 iterations and a quality control criterion of 0.0002, unless stated otherwise.

### Live-cell imaging of lysosomal tubulation

Cells were seeded into 4-well imaging chamber (Ibidi) one day prior to imaging to achieve a confluency of approximately 70% at the time of acquisition. Lysosomal tubulation and fission events were enriched by treating cells with 1.6 µM YM201636 in a complete DMEM for 2 h^10^. Cells were washed with imaging buffer 3 times and allowed to recover in imaging buffer. Images were acquired for 3 h at an interval of 15 min. Where specified, 100 nM Lysotracker Deep Red was added 20 min prior to the imaging to stain acidic compartments. Similarly, degradative compartments were labelled with Magic Red dye as per manufacturer’s instructions.

Fission events of lysosomal tubules were captured by imaging cells immediately after washout of the drug. Imaging was performed in imaging buffer at 37°C with 5% CO_2_. Each cell was imaged for 2 min at an interval of 2 seconds. To assess the effect of knock-outs or mutants on either tubulation or fission, cells were allowed to recover for 30 min after drug washout in imaging buffer and imaged.

Nigercin was also used to enlarge endocytic compartment to visualize tubules^18^. Briefly, cells were treated with 10 µM nigericin for 20 min, washed four times in imaging buffer, and the imaging chamber was mounted onto an automated microscope stage of inverted Axio Observer microscope (Zeiss). Several fields of view (FOV) were selected for imaging per condition. The microscope was programmed to image all FOVs sequentially and repeat the imaging at 15 min intervals for 2 h.

The formation sucrosomes and subsequent lysosomal reformation was performed as described previously^7^. Cells were fed with 30 mM sucrose for 24 h and subsequently incubated with 0.5 mg/mL invertase in DMEM for 1 h. After the incubation, cells were imaged in imaging buffer at 37°C with 5% CO_2_ for 2 h at an interval of 15 min.

### Imaging of endolysosome during Ca^+2^ transport inhibition

HeLa cells transiently expressing LAMP1-GFP were seeded onto imaging chambers at a density of 7.5 × 10⁴ cells per well and cultured for 24 h prior to imaging to achieve optimal confluency. Cells were treated with 1.6 μM YM201636 together with 10 μM ML-SI3 (a TRPML1 inhibitor) for 2 h. Following treatment, cells were washed with imaging buffer and, as described in Fig. 8B, allowed to recover for 30 min either in drug-free medium or in medium containing 10 μM ML-SI3. Live-cell imaging was performed at 37°C in 5% CO_2_ using an inverted Axio Observer microscope (Zeiss) equipped with a Plan-Apochromat N 63×/1.40 oil DIC M27 objective, a Photometrics Prime 95B camera, and Zen 2.6 acquisition software. For the recovery experiments, cells maintained in ML-SI3 containing medium were washed after 30 min of imaging and subsequently incubated in drug-free medium for an additional 30 min. Imaging was continued during this period at 15-min intervals to visualize the resumption of endolysosomal reformation (ELR). In parallel experiments, mitochondrial Ca²⁺ import was inhibited using VDAC1 or VDAC2/3 inhibitors. Cells were treated with 1.6 μM YM201636 together with either 10 μM erastin or 10 μM VBIT-4 for 2 h and imaged under identical conditions as described for ML-SI3 treatment.

### Image analysis and quantification

The total number of lysosomes and LysoTracker-positive puncta were quantified using the *Analyze Particles* function in FIJI (ImageJ). Prior to quantification, images were deconvolved using Huygens Remote Manager (version 3.10.0, Scientific Volume Imaging). Otsu’s thresholding method was applied to define signal boundaries, and total particle counts were measured per cell across eight time points. The frequency of tubulation, budding, or shape distortion events was quantified manually by visual inspection of the first 100 events in a total of 30 cells, each imaged for 2 min.

To visualize and quantify the distribution and intensity of lysosomal luminal markers, individual tubulating endolysosomes were imaged. A line was drawn through the lumen and tubules of LAMP1-positive (green) endolysosomes, and the intensity of the magenta channel (LysoTracker, Magic Red, Cathepsin D-RFP, or EGFR-mCherry) was measured in FIJI. Intensity profiles were plotted as gray value versus distance using GraphPad Prism 10.

The average lysosomal area was measured using the *Analyze Particles* function in FIJI within 100 μm² regions of interest (ROIs). Otsu’s thresholding was applied to define lysosomal structures. Area values from 90 ROIs across 30 cells from three independent experiments were analyzed and plotted in GraphPad Prism 10. For quantification of lysosomal tubule length, deconvolved images were analyzed in FIJI. Tubules were defined as particles with circularity between 0 and 0.5. Tubule length was measured by drawing a line along the tubular signal, and values were expressed in µm. At least 90 tubules from 30 cells across three independent experiments were quantified and plotted in GraphPad Prism 10.

For colocalization analysis of mCherry-DRP1 and LAMP1-EGFP in HeLa cells, wide-field fluorescence images were deconvolved using Huygens Remote Manager (version 3.10.0). Deconvolution was performed using a theoretical point spread function (PSF), automatic aberration correction, and the classic maximum likelihood estimation algorithm with 50 iterations, a quality factor of 0.0002, acuity mode set to 50, and automatic background estimation. Rectangular ROIs were drawn on deconvolved images, and Mander’s colocalization coefficients were calculated using the JACoP plugin in FIJI. ROI-based values were plotted using GraphPad Prism 10.

Mean fluorescence intensities of LysoTracker, Magic Red, and filipin (for cholesterol accumulation) were quantified per cell in DRP1 knockout cells using the *Analyze Particles* function in FIJI. Mean fluorescence values from individual cells (n = 30, from three independent experiments) were plotted as individual data points in GraphPad Prism 10. The spatial distribution of LysoTracker and Magic Red puncta was determined by drawing a line from the nucleus (Hoechst signal) to the cell periphery (actin mask). Intensity histograms were extracted in FIJI and plotted as gray value versus distance (μm) using GraphPad Prism 10.

To assess the proximity of DRP1, ER, and mitochondria to tubulating endolysosomes, super-resolution images were deconvolved using Huygens Remote Manager (version 3.10.0). Individual contact events were identified, and a line was drawn across each site. Intensity profiles along the line were extracted in FIJI and plotted as gray value versus distance (μm) in GraphPad Prism 10. The percentage occurrence of ER and mitochondrial contacts at fission sites was quantified manually.

Mitochondrial calcium signals were quantified by measuring the mean fluorescence intensity of Mito-RCaMP1h in individual cells at three time points (15-min intervals) using FIJI. Data from 30 cells across three independent experiments were plotted as individual values in GraphPad Prism 10.

All image panels were assembled using the OMERO image data management system.

### Immunobloting

Cells were lysed 24 h after seeding in lysis buffer (1% Triton X-100, 150 mM NaCl, 20 mM Tris-HCl pH 7.5, 1 mM EDTA, 1 mM EGTA, and Halt protease inhibitor cocktail). Lysates were denatured in Laemmli buffer at 65°C for 10 min, resolved on 10% SDS–PAGE, and transferred onto nitrocellulose membranes (Amersham). Membranes were blocked for 30 min in TBST (20 mM Tris-HCl pH 7.6, 150 mM NaCl, 0.1% Tween 20) containing 5% (w/v) non-fat dry milk and incubated overnight at 4°C with primary antibodies. After washing, membranes were incubated for 2 h at room temperature with HRP-conjugated secondary antibodies. Signals were detected using WesternBright ECL substrate (cat. K-12045-D50/ Advansta) and imaged with a Fusion FX imager (Vilber Lourmat). Band intensities were quantified in Fiji.

### Proximity biotinylation and Immuno Pulldown

HeLa cells were seeded in 6-well plates at a density of 5 × 10⁵ cells per well in triplicate. Two wells per condition (1 × 10⁶ cells) were used for each experiment. Cells were transfected with 1.5 μg of the LAMP1-TiD construct per well 24 h prior to biotinylation. Where indicated, cells were treated with 1.6 μM YM201636 for 2 h, followed by a 30-min washout and recovery in drug-free medium. During the recovery period, 50 μM biotin was added to the culture medium to enable biotinylation of proximal proteins. The reaction was quenched by transferring the cells to ice and washing them five times with ice-cold PBS. Cells were lysed in RIPA buffer (50 mM Tris-HCl pH 7.5, 150 mM NaCl, 0.1% (w/v) SDS, 0.5% (w/v) sodium deoxycholate, and 1% (v/v) Triton X-100) supplemented with protease inhibitors. Lysates were clarified by centrifugation at 12,000 rpm for 10 min at 4°C, and the supernatant was collected.

For streptavidin pulldown, 300 μg of total protein from each sample was incubated with 50 μL of streptavidin-conjugated beads in 500 μL of RIPA buffer for at least 1 h at 4°C. Beads were washed twice with RIPA buffer (1 mL, 2 min at room temperature), once with 1 M KCl (1 mL, 2 min), once with 0.1 M Na2CO3 (1 mL, ∼10 s), once with 2 M urea in 10 mM Tris-HCl pH 8.0 (1 mL, ∼10 s), and twice more with RIPA buffer (1 mL, 2 min each).

### Digestion of TurboID samples and proteomics analysis

The resin was resuspended in elution buffer (5% SDS, 10 mM TCEP, 0.1 M TEAB, pH 8.0), incubated for 10 min at 95°C, cleared by centrifugation and eluate was transferred into a new tube. Proteins were alkylated in 20 mM iodoacetamide for 30 min at 25°C and digested using S-Trap™ micro spin columns (Protifi) according to the manufacturer’s instructions. Shortly, 12 % phosphoric acid was added to each sample (final concentration of phosphoric acid 1.2%) followed by the addition of S-trap buffer (90% methanol, 100 mM TEAB pH 7.1) at a ratio of 6:1. Samples were mixed by vortexing and loaded onto S-trap columns by centrifugation at 4,000 g for 1 min followed by three washes with S-trap buffer. Digestion buffer (50 mM TEAB pH 8.0) containing sequencing-grade modified trypsin (1/25, w/w; Promega, Madison, Wisconsin) was added to the S-trap column and incubated for 1h at 47°C. Peptides were eluted by the consecutive addition and collection by centrifugation at 4,000 g for 1 min of 40 µl digestion buffer, 40 µL of 0.2% formic acid and finally 35 µL 50% acetonitrile, 0.2% formic acid. Samples were dried under vacuum and stored at −20 °C until further use.

Dried peptides were resuspended in 0.1% aqueous formic acid and subjected to LC-MS/MS analysis using a Orbitrap Fusion Lumos Mass Spectrometer fitted with an EASY-nLC 1200 (both Thermo Fisher Scientific) and a custom-made column heater set to 60°C. Peptides were resolved using a RP-HPLC column (75um × 30cm) packed in-house with C18 resin (ReproSil-Pur C18–AQ, 1.9 µm resin; Dr. Maisch GmbH) at a flow rate of 0.2 µL/min. The following gradient was used for peptide separation: from 5% B to 12% B over 5 min to 35% B over 40 min to 50% B over 15 min to 95% B over 2 min followed by 18 min at 95% B. Buffer A was 0.1% formic acid in water and buffer B was 80% acetonitrile, 0.1% formic acid in water.

The mass spectrometer was operated in DDA mode with a cycle time of 3 seconds between master scans. Each master scan was acquired in the Orbitrap at a resolution of 120,000 FWHM (at 200 m/z) and a scan range from 375 to 1600 m/z followed by MS2 scans of the most intense precursors in the linear ion trap at “Rapid” scan rate with isolation width of the quadrupole set to 1.4 m/z. Maximum ion injection time was set to 50 ms (MS1) and 35 ms (MS2) with an AGC target set to 1e6 and 1e4, respectively. Only peptides with charge state 2 - 5 were included in the analysis. Monoisotopic precursor selection (MIPS) was set to Peptide, and the Intensity Threshold was set to 5e3. Peptides were fragmented by HCD (Higher-energy collisional dissociation) with collision energy set to 35%, and one microscan was acquired for each spectrum. The dynamic exclusion duration was set to 30 s.

The acquired raw-files were searched using MSFragger (v. 4.1) implemented in FragPipe (v. 22.0) against a Homo sapiens database (consisting of 20360 protein sequences downloaded from Uniprot on 20220222) and 392 commonly observed contaminants using the “LFQ-MBR” workflow. Quantitative data was exported from FragPipe and analyzed using the MSstats R package v.4.13.0. (https://doi.org/10.1093/bioinformatics/btu305). Data was not normalised, imputed using “AFT model-based imputation” and p-values / q-values for pairwise comparisons were calculated using the limma package (DOI: 10.18129/B9.bioc.limma).

### Statistics

Statistical analyses were performed using Prism (GraphPad Software LLC). The statistical tests used and corresponding *P* values are indicated in the figure legends. Data sets were tested for normality or lognormality using the Anderson-Darling, Shapiro-Wilk, Kolmogorov-Smirnov, and D’Agostino-Pearson tests. Differences in standard deviation were assessed using the Brown-Forsythe and Bartlett’s tests, and differences in variance were evaluated using the *F* test. A significance level of 0.05 was applied to all tests. Based on data distribution and variance, the appropriate statistical test was selected; for example, if data were not normally distributed and multiple comparisons were required, nonparametric tests such as the Kruskal-Wallis test followed by Dunn’s multiple-comparisons test were used.

## Acknowledgments

We wish to thank. M. Spiess, M. Ryan, A.M. Lennon Dumenil and O. Chakrabarty for cell lines and plasmids. We acknowledge the expert help from Imaging Core facility (IMCF) and the FACS facility of the Biozentrum of the University of Basel. This work was supported by the University of Basel and the Swiss National Science Foundations (320030-231859, 310030-197779) to AS.

## Data availability

All data are included in the manuscript, except for the proteomics data, which have been submitted to the ProteomeXchange Consortium (https://www.proteomexchange.org/) via the MassIVE partner repository (https://massive.ucsd.edu/) with MassIVE data set identifier MSV000100521 and ProteomeXchange identifier PXD073275.

## Declaration of conflict of interest

The authors declare no conflict of interest.

---

**Figure S1.**
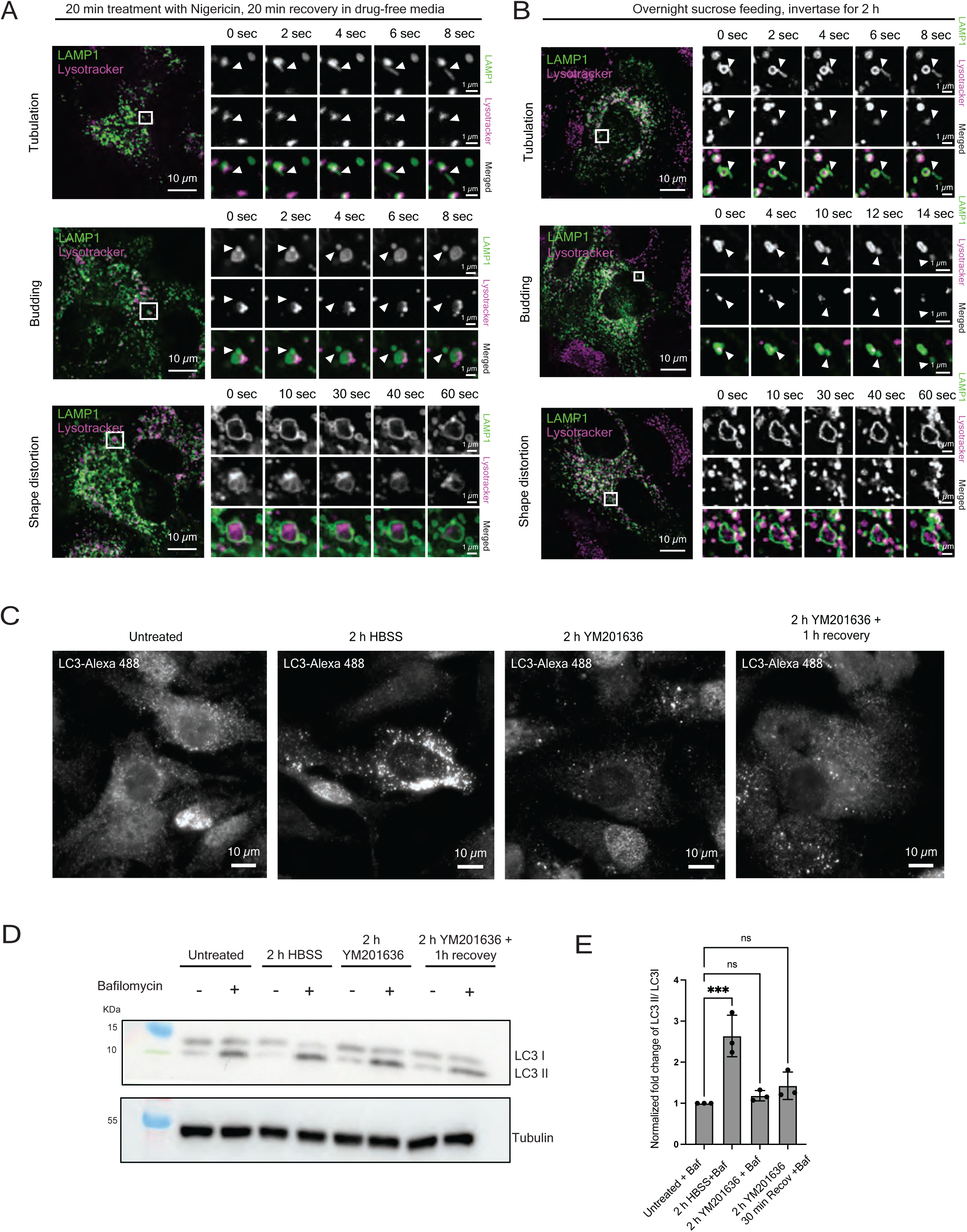
YM201636 treatment as a tool to synchronize ELR. (A) Live-cell time-lapse imaging of cells expressing LAMP1-GFP (green) and stained with Lysotracker Deep Red (magenta). Cells were treated with Nigericin to enlarge endocytic compartments. Nigericin was washed out and imaging was performed after 20 min of recovery. Each cell was imaged for 2 min at an interval of 2 sec. (B) Live-cell time-lapse imaging of cells expressing LAMP1-GFP (green) and stained with Lysotracker Deep Red (magenta) treated with sucrose and invertase. Cells were fed with sucrose overnight and treated with invertase for 2 h before imaging to initiate ELR. Each cell was imaged for 2 min at an interval of 2 sec. (C) Immunofluorescent image of cells stained for LC3. Cells treated with YM20163 and recovery conditioned were fixed using 4%PFA while keeping starved and untreated cells as positive and negative control respectively. After fixation, coverslips were stained with anti-LC3 antibodies for imaging. (D) Immunoblot analysis to check the autophagic flux in cells treated with YM2016136 and recovery condition. Starvation with HBSS was used as a positive control for LC3 II lipidation. Tubulin was used as a loading control. (E) Fold change of LC3 II/LC3 I was assessed and plotted for bafilomycin positive conditions. Plot represents the mean fold change across n=3 biological replicates; One way ANOVA with Dunn’s multiple comparison; ***P= 0.0005 (untreated vs HBSS), ns P = 0.8087 (untreated vs 2 h YM201636), ns P = 0.2845 untreated vs 2 h YM201636 + 1h recovery)

**Figure S2.**
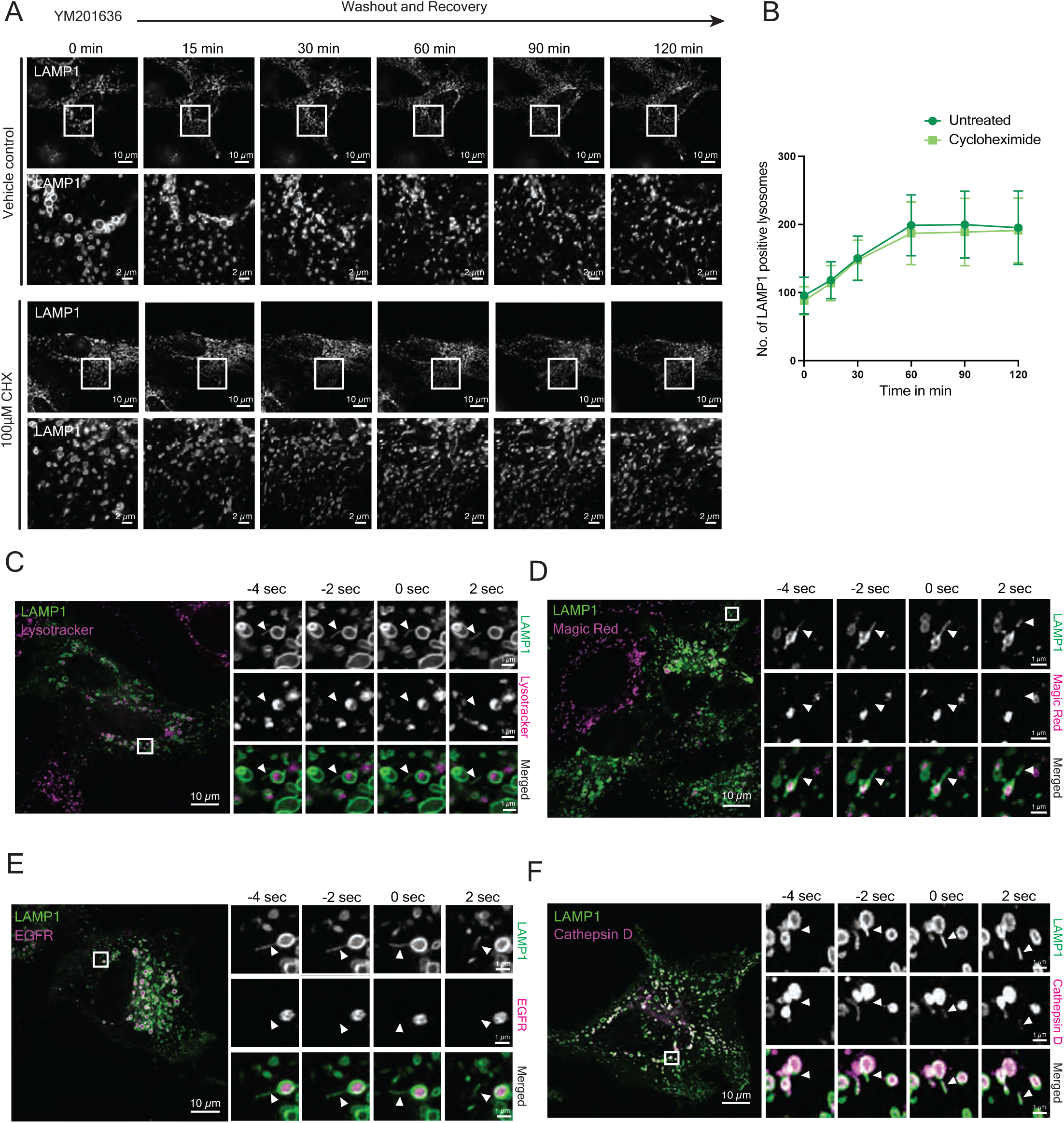
Active translation is dispensable for repopulation of lysosomes during ELR. (A) Live-cell time-lapse imaging of cells expressing LAMP-GFP to assess the effect of translation inhibition on ELR kinetics. Cells were treated with YM201636 for 2 h followed by washout and recovery to initiate ELR. During recovery, cycloheximide was introduced in the recovery media while keeping media with ethanol as a vehicle control. Cells were imaged for 2 h at an interval of 15 min. (B) Quantification of LAMP1-positive structures from (A). The line plot shows the number of LAMP1-positive compartments per cell over the recovery period. Data represent measurements from 30 cells. The central line indicates the mean; error bars represent the standard deviation (SD). C) Live-cell super resolution imaging of tubulating endolysosome from a cell expressing LAMP1-GFP (green) and stained with Lysotracker Deep Red (magenta). Cells were imaged for 2 min at an interval of 2 sec. Enlarged ROI shows the fission events of LAMP1-positive compartments (green) and corresponding LysoTracker signal (magenta). (D) Live-cell super confocal imaging of tubulating endolysosome from a cell expressing LAMP1-GFP (green) and stained with Magic Red (magenta). Cells were imaged for 2 min at an interval of 2 sec. Enlarged ROI shows the fission events of LAMP1-positive compartments (green) and corresponding Magic Red signal (magenta). E) Live-cell super resolution imaging of tubulating endolysosome from a cell co-expressing LAMP1-GFP (green) and EGFR-mCherry. Cells were stimulated with EGF for 30 min to increase EGFR uptake and imaged for 2 min at an interval of 2 sec. Enlarged ROI shows the fission events of LAMP1 positive compartments (green) and corresponding EGFR-mCherry signal (magenta). (F) Live-cell super confocal imaging of tubulating endolysosome from a cell co-expressing LAMP1-GFP (green) and cathepsin D-RFP (magenta). Cells were imaged for 2 min at an interval of 2 sec. Enlarged ROI shows the fission events of LAMP1-positive compartments (green) and corresponding cathepsin D signal (magenta).

**Figure S3.**
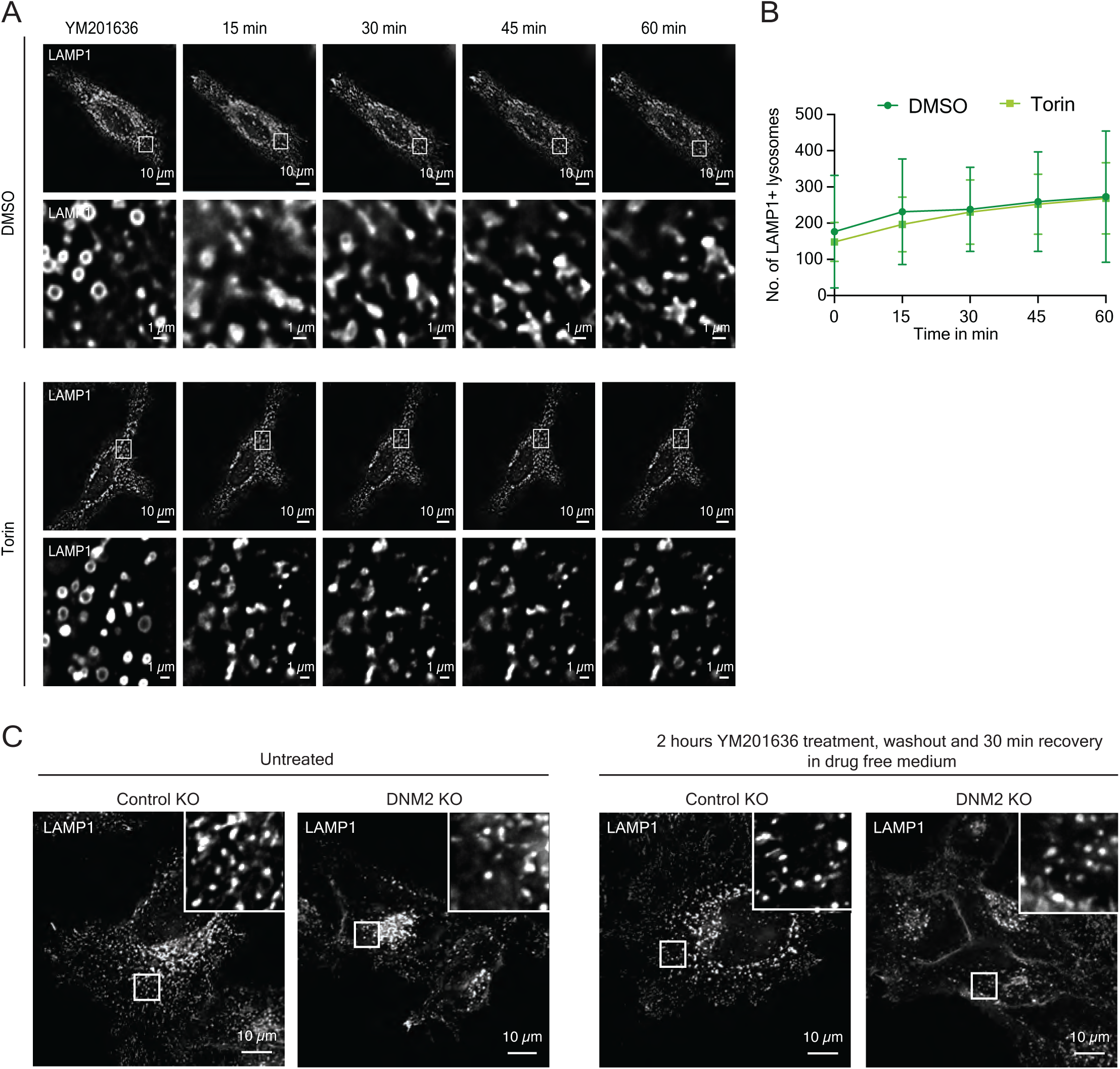
ELR proceeds without mTOR or dynamin-2. (A) Live-cell time-lapse imaging of cells expressing LAMP1–GFP to assess the effect of mTOR inhibition on ELR kinetics. Cells were treated with YM201636 for 2 h followed by washout and recovery to initiate ELR. During recovery, the mTOR inhibitor torin was added to the medium, with DMSO as vehicle control. Cells were imaged for 1 h at 15-min intervals. (B) Quantification of LAMP1-positive structures from (A). Line plot shows the number of LAMP1-positive compartments per cell over the recovery period. Data represent measurements from 30 cells. The central line indicates the mean; error bars indicate the standard deviation**. (**C) Live-cell imaging of control knockout and dynamin-2 knockout cells expressing LAMP1-GFP to assess the role of dynamin 2 in endolysosomal tubule fission. Cells were imaged after 2 h YM201636 treatment followed by 30-min washout and recovery. Untreated cells were used as controls.

**Figure S4.**
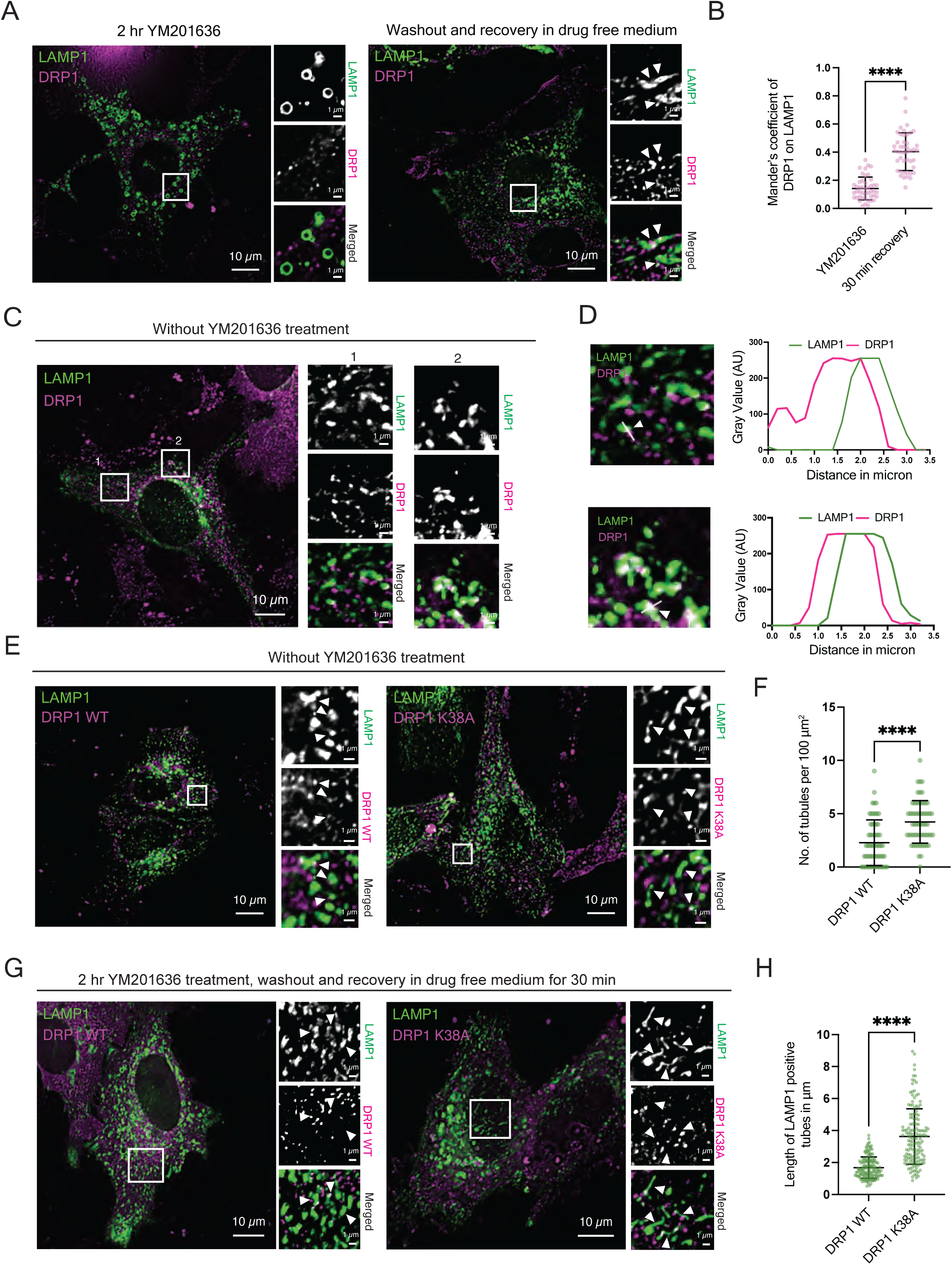
The GTPase activity of DRP1 is indispensable for endolysosomal tubule fission. (A) Live-cell imaging of cells co transfected with LAMP1-GFP (green) and DRP1-mCherry (magenta) to assess the localization of DRP1 on lysosomes during reformation. Cells were imaged after treating with YM201636 for 2 h to inhibit the ELR and also after washout and 30 min of recovery to visualize the DRP1 on tubulating structures. (B) Co-localization of DRP1 with LAMP1 was measured in both YM201636 and recovery condition using Mander’s coefficient of DRP1 on LAMP1. Each dot represents the average Mander’s coefficient value of 100 μm^2^ ROI. The plot shows mean Mander’s coefficient of 45 ROIs obtained from 15 cells in n=3 biological replicates; Unpaired *t*-test: ****P < 0.1 ×10^−14^ (YM201636 vs 30 min recovery). (C) Live-cell imaging of cells co transfected with LAMP1-GFP (green) and DRP1-mCherry (magenta) to assess the localization of DRP1 on lysosomes in untreated condition. (D) DRP1 (magenta) localized on the LAMP1-positive compartments (green) are highlighted in enlarged ROI using white arrowheads. The localization was also plotted using line plots measuring the gray values of LAMP1-GFP (green) and DRP1-mCherry (magenta) channels over the distance in µm. (E) Live-cell imaging of cells co-expressing LAMP-GFP and wild-type DRP1 or the DRP1 K38A mutant. (F) Quantification of (E). Number of tubular lysosomes were quantified and plotted. Each dot represents the number of LAMP1-positive tubules per 100 μm^2^ ROI. The plot shows the mean number of tubular structures of 90 ROIs measure across 30 cells in n=3 replicate; Unpaired *t*-test: ****P = 0.4659 ×10^−6^ (DRP1 WT vs DRP1 K38A). (G) Live-cell imaging of cells co-expressing LAMP-GFP and wild type DRP1 or the DRP1 K38A mutant. Cells were imaged after 2 h YM201636 treatment followed by washout and recovery of 30 min. (H) Quantification of LAMP1-positive tubule length from (G). Each data point represents the length of a single tubule. The plot shows the mean length of tubules from 30 cells across n=3 biological replicates; Unpaired *t*-test: ****P < 0.1 ×10^−14^ (DRP1 WT vs DRP1 K38A).

**Figure S5.**
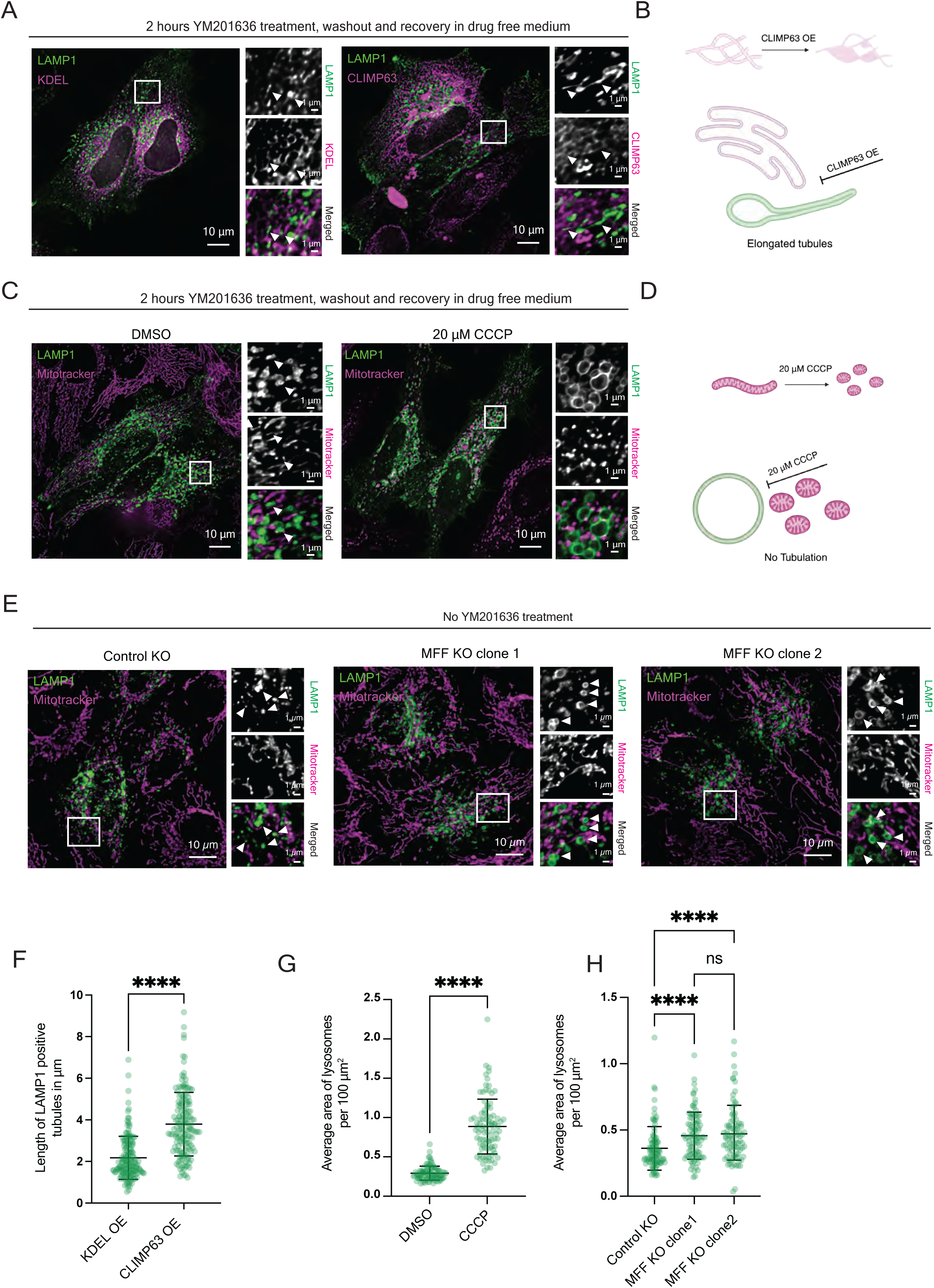
Contact between ER, mitochondria and endolysosome drive tubules fission during ELR. (A) Disruption of ER contact by overexpressing CLIMP63. Cells were transfected with LAMP1-GFP (green) and KDEL-mCherry or CLIMP63-mCherry. Imaging was performed after 2 h of YM201636 treatment, washout, and 30 min of recovery. White arrows depict the LAMP1-positive compartment with respective ER morphology. (B) Schematic representation of CLIMP63 overexpression and its implication on ELR. (C) Mitochondrial morphology was disturbed by treating cells with 20 μM CCCP for 30 min. Cells expressing LAMP1-GFP (green) were treated with YM201636 for two h followed by washout and 30 min recovery. Twenty μM CCCP was introduced in the recovery media. Mitochondria were stained using MitoTracker Deep Red (magenta). Images were taken after 30 min of recovery to observe the effect. White arrow shows the morphology of the LAMP1-positive compartment in DMSO versus CCCP-treated cells. (D) Schematic representation of CCCP treatment on mitochondria and its implication on ELR. (E) Live-cell imaging of LAMP1-positive lysosomes (green) in control KO and MFF knockout (KO) cells. Cells were imaged after staining with MitoTracker Deep Red (magenta). White arrowheads highlight the LAMP-1positive compartments in the ROIs. (F) Quantification of tubules length of endolysosomes from data (A). Each dot represents the length of individual tubules and the plot shows the mean of 90 tubules measured in 30 cells across n=3 biological replicates; Unpaired *t*-test: ****P < 0.1 ×10^−14^ (KDEL OE vs CLIMP OE). (G) Quantification of the average area of LAMP1-positive structures for (C). Each dot represents the average area per 100 µm² ROI. The plot shows the mean area from a total of 90 ROIs from 30 cells across n= 3 biological replicates; Unpaired *t*-test: ****P < 0.1 ×10^−14^ (DMSO vs CCCP). (H) Quantification of the average area of LAMP1-positive structures for (E). Each dot represents the average area per 100 µm² ROI. The plot shows the mean area from a total of 90 ROIs from 30 cells across n = 3 biological replicates; Kruskal-Wallis test with Dunn’s multiple comparisons: ****P = 0.5356 ×10^−4^ (Control KO vs MFF KO C1), ****P = 0.2273 ×10^−4^ (Control KO vs MFF KO C 2), ns P > 0.9 ×10^−14^ (MFF KO C1 vs MFF KO C2).

**Movie S1. Reformation of lysosomes after YM201636 washout**

Time-lapse imaging of lysosome reformation following YM201636 treatment and washout. LAMP1-GFP expressing cells were stained with LysoTracker Deep Red and imaged every 15 min after washout. Three representative examples of cells undergoing endolysosomal reformation (ELR) are shown. See Fig.1

**Movie S2. ELR tubules are positive for cathepsin D**

Time-lapse imaging of cells expressing LAMP1-GFP and CTSD-RFP during recovery from YM201636 treatment. Three representative tubulation and fission events show cathepsin D signal in both the globular and tubular regions of reforming endolysosomes, with fissioned tubules remaining positive for cathepsin D. See Fig. 2

**Movie S3. DRP1 at the fission site**

Time-lapse imaging of cells expressing LAMP1-GFP and DRP1-mCherry during recovery from YM201636 treatment. Three representative examples show endolysosomal tubule fission events with DRP1 localized at the fission site. See Fig. 4

**Movie S4. Inability of DRP1 K38A mutant to induce scission**

Time-lapse imaging of cells expressing LAMP1-GFP and DRP1 K38A-mCherry during recovery from YM201636 treatment. Three representative examples show recruitment of the DRP1 (K38A) mutant to endolysosomal tubules without subsequent membrane fission. See Fig. S4.

## Notes

### Competing Interest Statement

The authors have declared no competing interest.

